# Mapping memories: pulse-chase labeling reveals AMPA receptor dynamics during memory formation

**DOI:** 10.1101/2023.05.26.541296

**Authors:** Doyeon Kim, Pojeong Park, Xiuyuan Li, J. David Wong Campos, He Tian, Eric M. Moult, Jonathan B. Grimm, Luke Lavis, Adam E. Cohen

## Abstract

A tool to map changes in synaptic strength during a defined time window could provide powerful insights into the mechanisms governing learning and memory. We developed a technique, Extracellular Protein Surface Labeling in Neurons (EPSILON), to map α-amino-3-hydroxy-5-methyl-4-isoxazolepropionic acid receptor (AMPAR) insertion *in vivo* by pulse-chase labeling of surface AMPARs with membrane-impermeable dyes. This approach allows for single-synapse resolution maps of plasticity in genetically targeted neurons during memory formation. We investigated the relationship between synapse-level and cell-level memory encodings by mapping synaptic plasticity and cFos expression in hippocampal CA1 pyramidal cells upon contextual fear conditioning (CFC). We observed a strong correlation between synaptic plasticity and cFos expression, suggesting a synaptic mechanism for the association of cFos expression with memory engrams. The EPSILON technique is a useful tool for mapping synaptic plasticity and may be extended to investigate trafficking of other transmembrane proteins.

## Introduction

Changes in synaptic strength are an important component of learning and memory ^1–6^, but the rules which map a memory onto a specific set of synapses are not well understood. Which synapses represent which memories? How are changes in synaptic strength related to other markers of memory, such as expression of immediate early genes? To answer these questions, one would like a tool to map changes in synaptic strength during a defined time window in genetically defined neurons.

The synaptic density of α-amino-3-hydroxy-5-methyl-4-isoxazolepropionic acid receptors (AMPARs) is a major contributor to synaptic strength. The density of AMPARs changes during long-term potentiation (LTP) and long-term depression (LTD) ^7–9^. AMPARs are stored in intracellular vesicles, which fuse with the postsynaptic membrane during LTP, exposing the N-terminal glutamate-binding domain to the extracellular space. The lumen of the intracellular vesicles is acidic, so AMPAR insertion can be monitored via fluorescence of a fusion of a pH-sensitive fluorescent protein, such as Super Ecliptic pHluorin (SEP), to the N terminus of the AMPAR subunit GluA1. *In vivo* 2-photon microscopy of SEP-GluA1 has been used to observe AMPAR insertion at individual synapses during memory formation ^10–17^. However, despite progress in machine learning-based image analysis,^18^ the requirement for high resolution imaging *in vivo* restricts application of this technique to superficial cortex.^19^ Further, the requirement for head-fixed imaging constrains the possible behavioral paradigms ^20^. The technique of GFP reconstitution across synaptic partners (GRASP)^21,22^ and its enhancement (eGRASP)^23^ allow mapping of synaptic connections between defined neuronal populations, including under control of activity dependent promoters. These techniques separate the *in vivo* recording from the *ex vivo* measurement and so are applicable to deep brain regions, but do not directly probe the strength or timing of plasticity events.

Pulse-chase labeling of proteins with HaloTag-ligand dyes has been used to probe protein turnover at the level of translation and degradation *in vivo*, via a technique called DELTA (Dye Estimation of the Lifetime of proTeins in the brAin) ^24^. Since the dyes used in DELTA were membrane permeable, this technique was not sensitive to fine details of membrane trafficking, such as whether the protein was inside or on the surface of the cell.

Here we developed an approach to map AMPAR insertion by pulse-chase labeling of surface AMPARs with membrane-impermeable fluorescent dyes. We call the technique EPSILON (Extracellular Protein Surface Labeling in Neurons). We fused the self-labeling HaloTag (HT) protein to the N terminus of GluA1 (Fig. 1A). We expressed HT-GluA1 in neurons, and then saturated surface-exposed HT-GluA1 via direct brain injection of a membrane-impermeable HaloTag-ligand (HTL) dye (Fig. 1B). A second dye of a different color was then added to label newly exposed HT-GluA1, and the animal was exposed to a variety of learning paradigms (Fig. 1C). Subsequent *ex vivo* multi-color imaging revealed maps of AMPAR exocytosis with single-synapse resolution across large volumes of brain tissue and in multiple brain regions.

**Fig. 1.**
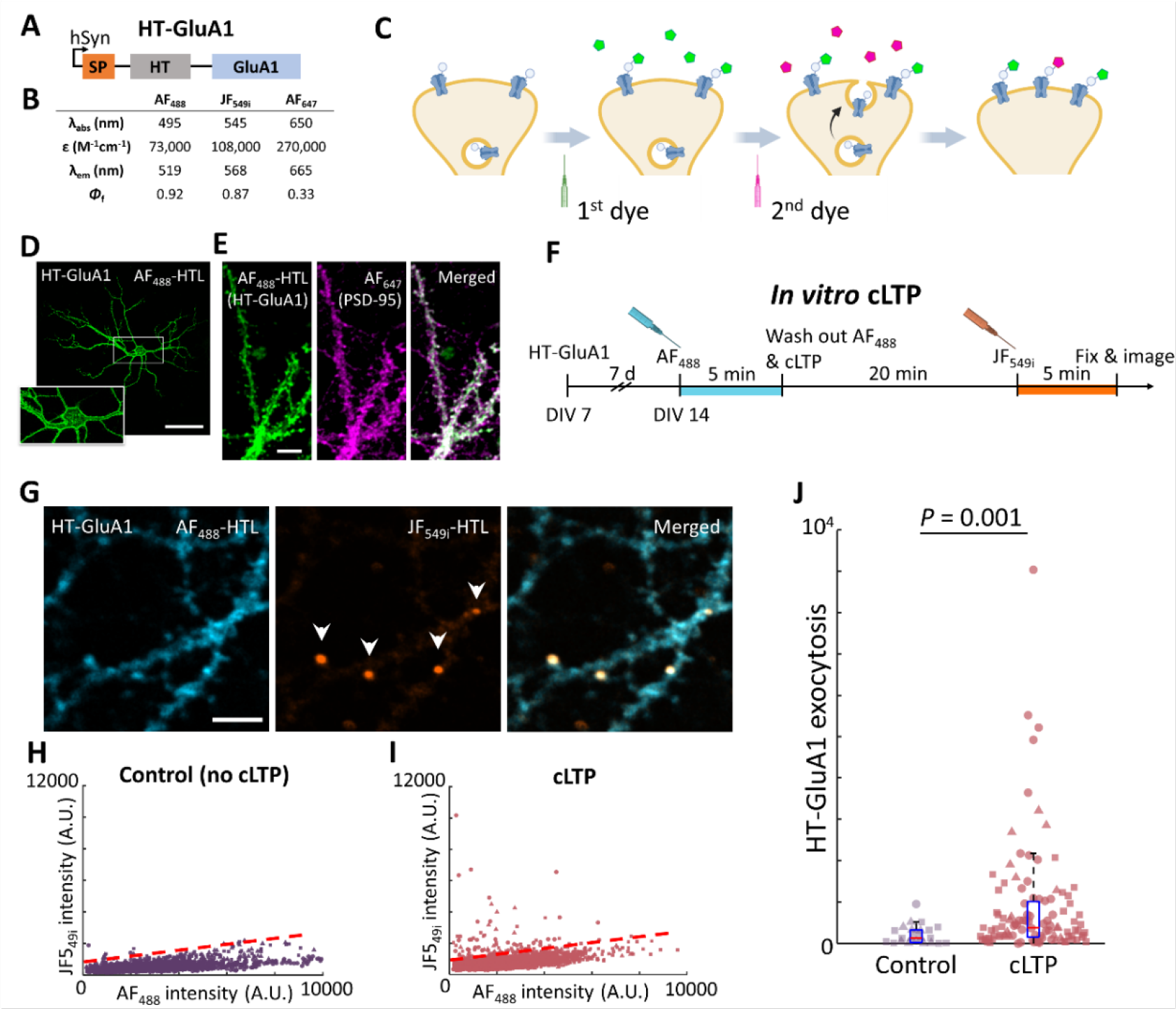
EPSILON tags freshly exposed AMPARs. **(A)** Composition of HT-GluA1. SP: GluA1 signaling peptide, HT: HaloTag receptor. **(B)** Membrane-impermeable Ligand (HTL) dyes used in this work. **(C)** EPSILON scheme for sequential labeling of surface-exposed AMPARs using membrane-impermeable HTL dyes. Preexisting surface HT-GluA1s are labeled with the first dye, followed by injection of the second dye to label newly exocytosed HT-GluA1. The two populations are analyzed using *ex vivo* multi-color imaging. **(D)** Cultured neuron expressing HT-GluA1. Surface HT-GluA1 was stained with AF_488_-HTL. Scale bar, 100 μm. **(E)** Postsynaptic trafficking of HT-GluA1 in a cultured neuron. (Left) HT-GluA1 stained with AF_488_-HTL. (Middle) PSD-95 (anti-PSD-95 immunostaining) from same region of interest. (Right) Merge. Scale bar, 10 μm. **(F)** Timeline for EPSILON pulse-chase labeling during chemical long-term potentiation (cLTP). DIV: days *in vitro*. **(G)** Images of spines labeled with two colors during cLTP. (Left) Preexisting HT-GluA1 stained with AF_488_-HTL before cLTP. (Middle) Newly exposed HT-GluA1 stained with JF_549i_-HTL after cLTP. Arrowheads indicate spines with high HT-GluA1 exocytosis. (Right) Merge. Scale bar, 10 μm. **(H), (I)** Scatterplots of spine fluorescence intensities in the two color channels for **(H)** control and **(I)** cLTP-treated neurons. Control: *n* = 2217 spines from 3 cultures; cLTP: *n* = 2513 spines from 3 cultures. Second dye intensity thresholds indicated with dashed lines (see materials and methods for calculation of threshold). Replicate dishes represented by different shape symbols. **(J)** Second dye intensity above threshold. Spines with second dye intensity higher than the defined threshold were included for analysis (control: *n* = 17 spines from 3 cultures; cLTP: *n* = 85 spines from 3 cultures). Replicate dishes represented by different shape symbols. Lower and upper bounds of the box plot: 25^th^ and 75^th^ percentiles, respectively; lower and upper whiskers: minimum and maximum, respectively; center lines: median. Two-sided Wilcoxon rank-sum test.

The hippocampus is necessary for formation and storage of spatial and contextual memories ^25,26^, but the physical nature of the hippocampal engram, or memory trace, remains unclear. On one hand, activation of subsets of hippocampal cells, termed engram neurons, is necessary and sufficient to recreate simple conditioned responses (e.g., freezing after contextual fear conditioning). These cells are often identified using expression of immediate early genes (e.g., cFos) ^27,28^. On the other hand, extensive research has shown that modulations of synaptic strength are also necessary for memory formation ^29^. A synaptic engram would allow partially overlapping subsets of neurons to be activated upon recall of different memories, a feat that would be difficult to achieve if memories were primarily stored at the level of whole-cell parameters such as gene expression. The relation between synapse-level and cell-level memory encodings is unclear ^30^.

We mapped synaptic plasticity and cFos expression in hippocampal CA1 pyramidal cells upon contextual fear conditioning (CFC). In mice subjected to CFC, but not in controls, we observed a tight correlation between the degree of synaptic plasticity and the cFos expression level. This finding suggests a synaptic mechanism for the observed association of cFos expression with memory engrams. We also observed more plasticity in perisomatic than in distal synapses; and clusters of plasticity among nearby synapses. These features may reflect interactions between plasticity and dendritic excitability properties ^31^. EPSILON tagging of AMPAR exocytosis is a powerful tool to investigate the distribution and time-course of synaptic plasticity, and we expect that the EPSILON approach could be applied to other transmembrane proteins.

## Main

### Development of EPSILON and validation in cultured neurons

We replaced the pH-sensitive SEP domain in SEP-GluA1 with HT to create HT-GluA1. This design used a flexible glycine linker between HT and GluA1 and retained the N-terminal GluA1 signal peptide^32^ to ensure proper protein trafficking.^10–17^

We first tested the EPSILON labeling scheme in cultured rat hippocampal neurons. We characterized (1) expression and trafficking of HT-GluA1, (2) the labeling kinetics of membrane impermeable HTL dyes, and (3) the turnover rate of surface HT-GluA1. HT-GluA1 showed good trafficking to the cell membrane (Fig. 1D and Fig. S1A). The colocalization of Alexa Fluor 488 (AF_488_)-labeled HT-GluA1 signal with the postsynaptic marker PSD-95 indicated that HT-GluA1 was correctly transported to the postsynaptic location (Fig. 1E and Fig. S1B). The labeling was saturated after approximately 300 seconds for AF_488_ (100 nM) and 100 seconds for JF_549i_ ^33^ (1 μM) (Fig. S2). In the subsequent pulse-chase labeling experiments in cultured neurons, we used conditions that saturated labeling (AF_488_: 100 nM for 5 min; JF_549i_: 1 μM for 5 min). We next determined the basal turnover rate of surface HT-GluA1 (Fig. S3) by pulsing with AF_488_, and then chasing with JF_549i_ after a variable delay. The ratio of the preexisting (labeled with AF_488_) and newly inserted (labeled with JF_549i_) surface HT-GluA1 yielded the turnover rate. The half-life of surface HT-GluA1 was approximately 30 min in cultured neurons.

We then asked whether HT-GluA1 could identify spines with high AMPAR exocytosis upon chemical LTP (cLTP) in cultured neurons (Fig. 1F). After saturating the surface HT-GluA1 with the first dye (AF_488_) and washing out unreacted dye, we induced cLTP by applying a cocktail of forskolin, rolipram, and picrotoxin (materials and methods) ^34,35^. We stained the cells with the second dye (JF_549i_) to label the freshly exocytosed HT-GluA1, and then fixed the cells and imaged each dye with a confocal microscope. We observed a significantly larger fraction of spines labeled with the second dye in cLTP-treated neurons compared to controls (Fig. 1G-I, cLTP: 85 of 2513 spines above threshold (3.4%), *n* = 32 neurons, 3 dishes; control: 17 of 2217 spines above threshold (0.8%), *n* = 28 neurons, 3 dishes; *P* = 0.009, two-sided Student’s t-test), consistent with prior results using the same cLTP cocktail ^35^. We also quantified the extent of AMPAR exocytosis in each spine by measuring the fluorescence of Dye 2 above baseline (materials and methods, Fig. 1J). Of spines labeled with Dye 2, the average Dye 2 fluorescence was higher in neurons treated with cLTP compared to controls (cLTP: 930 ± 150 counts, mean ± s.e.m., *n* = 32 neurons, 3 dishes; control: 320 ± 87 counts, mean ± s.e.m., *n* = 28 neurons, 3 dishes; *P* = 0.001, two-sided Wilcoxon rank-sum test). This analysis revealed that cLTP induced both an increase in the fraction of spines undergoing AMPAR exocytosis and also an increase in the amount of exocytosis in each spine.

### Validation of EPSILON in barrel cortex, *in vivo*

To characterize HT-GluA1 in the live mouse brain, we co-expressed HT-GluA1 and myc-GluA2 in layer 2/3 neurons in mouse barrel cortex via *in utero* electroporation (IUE) (Fig. 2A).^36^ As in prior experiments with SEP-GluA1, we used a 1:1 ratio of HT-GluA1 and myc-GluA2 to maintain the endogenous subunit stoichiometry.^10^ The HT-GluA1 formed bright puncta which localized in dendritic spines (Fig. S4). In some samples we co-expressed a morphological marker (membrane localized GFP) and verified that HT-GluA1 was well trafficked at spines (Fig. 2A). In acute brain slices, patch clamp recordings with extracellular stimulation of excitatory synaptic inputs showed that HT-GluA1/myc-GluA2 expression did not significantly alter the AMPAR-mediated EPSCs at various stimulus strengths or the AMPAR/NMDAR ratio (Fig. 2B-E). Current-clamp experiments confirmed that HT-GluA1/myc-GluA2 expression also did not significantly alter membrane resistance, membrane capacitance, resting potential, rheobase, or excitability (Fig. S5).

**Fig. 2.**
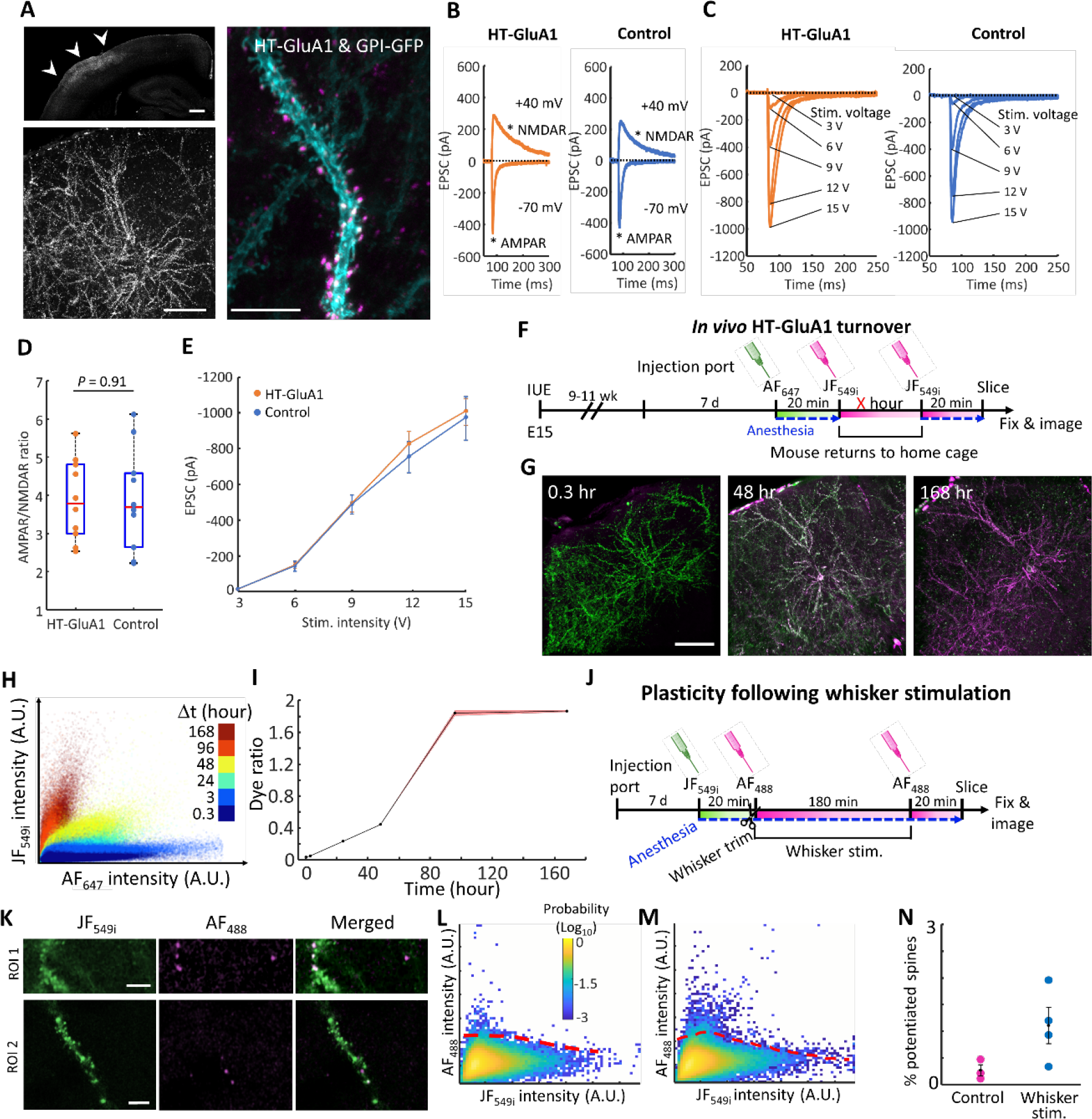
EPSILON reports sensory-induced AMPAR exocytosis *in vivo*. **(A)** Expression of HT-GluA1 in layer 2/3 pyramidal neurons in mouse barrel cortex. (Top left) Coronal section showing overall expression pattern. Arrowheads indicate barrel cortex. (Bottom left) Individual neurons expressing HT-GluA1. (Right) A dendrite co-expressing membrane-localized GFP (cyan) and HT-GluA1 (stained with AF_647_-HTL, magenta). Scale bars, 500 μm (top left), 200 μm (bottom left), and 10 μm (right). **(B)** AMPAR and NMDAR contributions to the EPSC evoked by an extracellular stimulating electrode. Data from layer 2/3 pyramidal neurons in barrel cortex, in acute brain slices. The AMPAR and NMDAR components (asterisks) were measured at holding potentials of -70 mV and +40 mV, respectively. **(C)** AMPAR-mediated EPSCs evoked with different stimulus intensities. **(D)** AMPAR/NMDAR ratio of HT-GluA1 expressing and control neurons (*n* = 10 neurons for each group). Two-sided Wilcoxon rank-sum test. **(E)** AMPAR-mediated EPSCs as a function of extracellular stimulus strength (*n* = 10 neurons for each group). Error bars show mean ± s.e.m. **(F)** EPSILON scheme for mapping turnover of surface AMPARs *in vivo*. E denotes embryonic age in days. **(G)** Pulse-chase labeled HT-GluA1 in layer 2/3 barrel cortex at different intervals after first chase; 0.3 h (left), 48 h (middle), and 168 h (right). Green: AF_647_, Magenta: JF_549i_. Scale bar, 200 μm. **(H)** Scatterplots of spine fluorescence intensities at different intervals after first chase (0.3 h: *n* = 312,246 spines from 3 mice; 3 h: *n* = 164,403 spines from 3 mice; 24 h: *n* = 458,181 spines from 3 mice; 48 h: *n* = 340,927 spines from 3 mice; 96 h: *n* = 26,765 spines from 3 mice; 168-hour: *n* = 121,503 spines from 3 mice). **(I)** Slopes of the scatterplots from (H) calculated by linear least-squares regressions. Error bars represent 95% confidence interval on the slope. **(J)** EPSILON scheme for mapping sensory-evoked AMPAR exocytosis. **(K)** Images of spines labeled with two colors upon whisker stimulation. (Top) Preexisting HT-GluA1 stained with JF_549i_-HTL before whisker stimulation. (Middle) Newly exocytosed HT-GluA1 stained with AF_488_-HTL during and after whisker stimulation. (Bottom) Merge. Scale bars, 5 μm. **(L), (M)** Density plots of spine fluorescence in **(L)** control and **(M)** whisker stimulated mice (control: *n* = 117,829 spines from 3 mice; whisker stimulated: *n* = 203,787 spines from 4 mice). Thresholds indicated with dashed lines (see materials and methods for calculation of threshold). **(N)** Fraction of spines exhibiting Dye 2 intensity above threshold. Each data point indicates one mouse (control: *n* = 117,829 spines from 3 mice; whisker stimulated: *n* = 203,787 spines from 4 mice). Error bars show mean ± s.e.m.

To quantify basal turnover of AMPARs (i.e. replacement of surface-exposed molecules with fresh ones from vesicles), we performed pulse-chase labeling of surface HT-GluA1 in the barrel cortex *in vivo* (Fig. 2F and Fig. S6, materials and methods). We saturated preexisting surface HT-GluA1 by intracortical injection of Dye 1 (1.4 μL of 1 μM AF_647_; see materials and methods for the synthesis of AF_647_-HTL). After 20 min—enough time for Dye 1 to react—we injected Dye 2 (1.4 μL of 10 μM JF_549_) at the same sites with ten-fold higher concentration to ensure that most new HT-GluA1 was labeled with Dye 2 and not residual Dye 1. The mice were returned to their home cage, except for those with an inter-injection delay of 0.3 hours, and after a variable delay, we injected Dye 2 again to ensure complete labeling of newly surface-exposed HT-GluA1. Confocal images of fixed brain sections mapped both dyes throughout 121,503-458,181 spines in 3 mice at each time-point (Fig. 2G), and the fluorescence intensities were quantified for each spine (Fig. 2H). The half-life for surface HT-GluA1 on spines *in vivo* was approximately 50 hours (Fig. 2I). The disparity in surface lifetime between culture (30 min) and *in vivo* (50 h) is consistent with prior reports which showed an acceleration of GluA1 protein turnover in slice cultures vs. *in vivo* ^37,38^. Pulse-chase experiments on shorter timescales could thus identify spines with accelerated AMPAR exocytosis. We also validated that the background autofluorescence and residual dyes contributed negligible fluorescence signal (Fig. S7A, materials and methods).

The adult barrel cortex exhibits synaptic plasticity in response to changes in sensory experience, such as stimulation or deprivation of a subset of whiskers ^10,39–41^. To tag spines with elevated GluA1 exocytosis upon the acute sensory stimulation, we performed EPSILON labeling while stimulating a subset of whiskers in an anesthetized mouse (Fig. 2J and Fig. S6A, materials and methods). Confocal images of L2/3 pyramidal neurons in fixed brain sections from the contralateral barrel cortex displayed spines with elevated Dye 2 (AF_488_), indicative of newly surface-exposed HT-GluA1 (Fig. 2K). Mice subjected to whisker stimulation had a larger fraction of spines with elevated Dye 2 compared to controls (whisker stimulated: 1.1 ± 0.34%, mean ± s.e.m. over 4 mice [n = 203,787 spines]; control: 0.27 ± 0.11%, mean ± s.e.m. over 3 mice [n = 117,829 spines]; Fig. 2, L-N, materials and methods). These findings demonstrate that pulse-chase HT-GluA1 AMPAR labeling can tag individual spines that undergo elevated AMPAR exocytosis *in vivo*.

### Relating hippocampal plasticity and cFos expression during fear conditioning

The CA1 region of the hippocampus is involved in formation and storage of context-related memories ^42^. Formation of cFos-expressing engram cells has been observed in this region ^27,28,43,44^, but studying *in vivo* AMPAR dynamics has been challenging due to the complex morphology of CA1 pyramidal neurons and their deeply embedded location ^45^. We used HT-GluA1 to investigate the relation between AMPAR dynamics and cFos expression in hippocampal CA1 pyramidal cells upon contextual fear conditioning (CFC).

We expressed HT-GluA/myc-GluA2 via IUE in hippocampal CA1 pyramidal neurons (materials and methods). As in the cortex, HT-GluA1 localized to dendritic spines (Fig. 3A and Fig. S4). We then used EPSILON tagging followed by fixed slice imaging to map newly exocytosed AMPARs in mice that underwent CFC, home cage control mice, and mice that were exposed to the context but not the conditioned stimulus (foot shock) (Fig. 3B, Fig. S6B and Fig. S8) ^46^. For each pyramidal neuron expressing HT-GluA1, we located every spine and quantified the fluorescence of Dye 1 and Dye 2 (materials and methods). We also quantified endogenous cFos in the nucleus via immunofluorescence in a third spectral channel (Fig. S9).

**Fig. 3.**
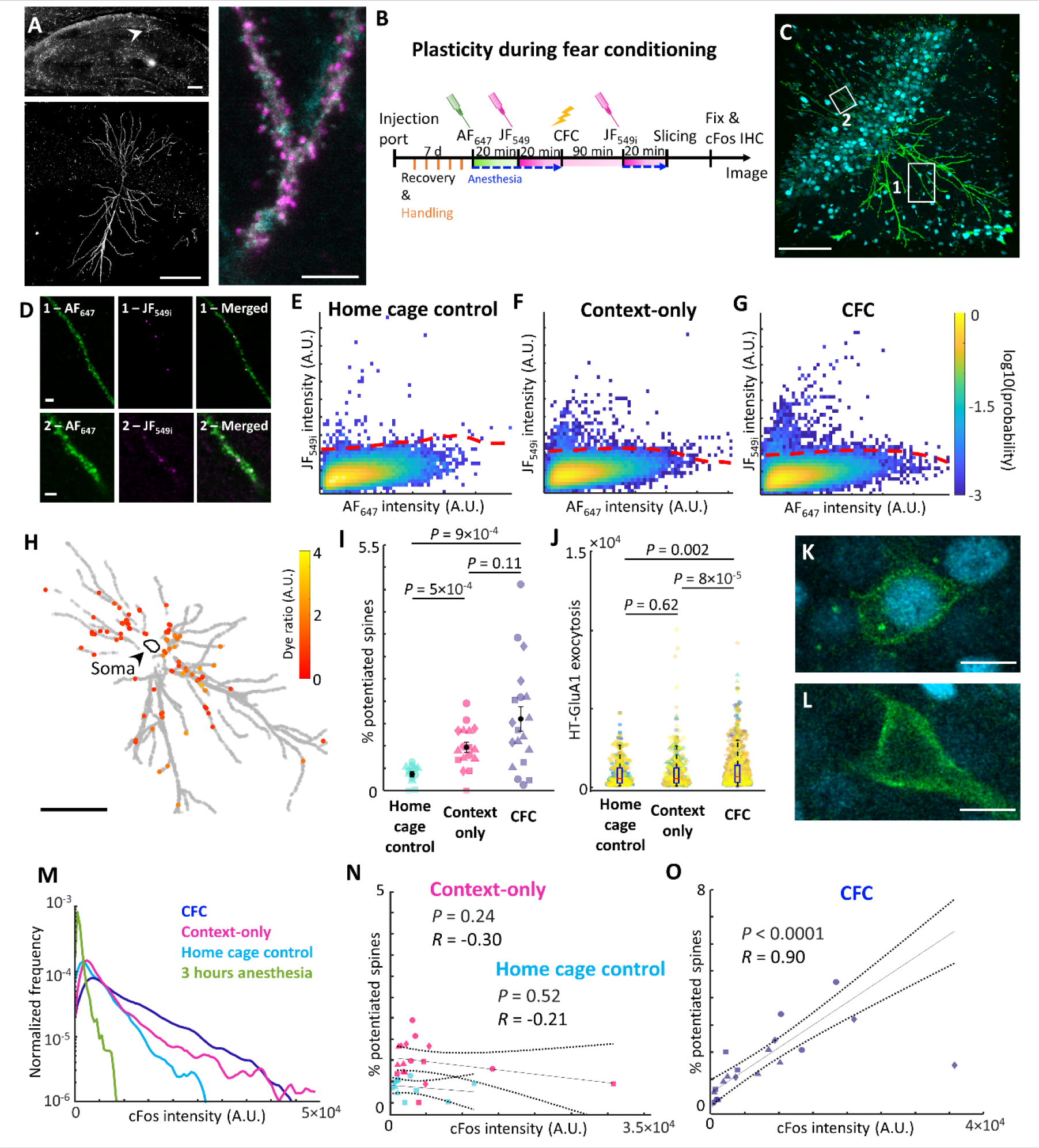
Contextual fear conditioning evokes correlated changes in AMPAR exocytosis and cFos expression. **(A)** Expression of HT-GluA1 in pyramidal neurons in mouse hippocampus CA1. (Top left) Coronal section showing overall expression pattern. Arrowhead indicates an expressing cell. (Bottom left) Single neuron expressing HT-GluA1. (Right) Dendrites co-expressing membrane-localized GFP (cyan) and HT-GluA1 stained with AF_647_-HTL, magenta. Scale bars, 200 μm (top left and bottom left), and 5 μm (right). **(B)** EPSILON scheme for tagging spines potentiated during CFC. IHC, immunohistochemistry. **(C)** Pyramidal neuron in CA1 after EPSILON tagging, CFC, and cFos immunostaining. Preexisting HT-GluA1 stained with AF_647_ (green) and immunostained cFos (cyan) are shown here. Scale bar, 200 μm. **(D)** Inset from C showing spines labeled with both dyes. (Left) Preexisting HT-GluA1 stained with AF_647_-HTL. (Middle) HT-GluA1 newly exposed during CFC, stained with JF_549i_-HTL. (Right) Merge. Scale bars, 10 μm. **(E)-(G)** Density plots of spine fluorescence in **(E)** home cage control, **(F)** context-only, and **(G)** CFC animals (home cage control: *n* = 43,659 spines from 3 mice; context-only: *n* = 74,801 spines from 4 mice; CFC: *n* = 75,200 spines from 4 mice). Thresholds indicated with dashed lines (see materials and methods for calculation of threshold). **(H)** Map of all spines (grey) and spines above Dye 2 threshold (color). The color scale indicates the fluorescence ratio, Dye 2/Dye 1. Scale bar, 200 μm. **(I)** Percentage of spines with Dye 2 above threshold (home cage control: *n* = 12 cells from 3 mice; context-only: *n* = 17 cells from 4 mice; CFC: *n* = 19 cells from 4 mice). Distinct mice indicated by different shape symbols. Error bars show mean ± s.e.m., two-sided Wilcoxon rank-sum test. **(J)** Dye 2 intensity above threshold in potentiated spines (home cage control: *n* = 215 spines from 12 cells from 3 mice; context-only: *n* = 683 spines from 16 cells from 4 mice; CFC: *n* = 1,016 spines from 19 cells from 4 mice). Distinct mice indicated with marker shape and distinct cells by marker color. Lower and upper bounds of the box plot: 25^th^ and 75^th^ percentiles; lower and upper whiskers: minimum and maximum; and center lines: median. Two-sided Wilcoxon rank-sum test. **(K), (L)** HT-GluA1 expressing CA1 neurons (green) with **(K)** high and **(L)** low cFos expression (cyan). Scale bars, 10 μm. **(M)** Distributions of cFos intensities in individual cells. Green: no exposure (*n* = 2,285 cells from 3 mice); cyan: home cage control (*n* = 1,922 cells from 3 mice); magenta: context-only (*n* = 5,074 cells from 4 mice); blue: CFC (*n* = 7,208 cells from 4 mice). **(N), (O)** Correlation between cFos intensity and the fraction of spines with high HT-GluA1 exocytosis from **(N)** home cage control (*n* = 12 cells from 3 mice) and context-only (*n* = 17 cells from 4 mice) and **(O)** CFC (*n* = 19 cells from 4 mice). Distinct mice indicated by different shape symbols. *R*, Pearson’s linear correlation coefficient, *P* value from two-sided Student’s t-test.

As in the barrel cortex, a subset of spines had elevated Dye 2 (JF_549i_) fluorescence, indicative of AMPAR exocytosis (Fig. 3C, D). We quantified the fraction of spines with elevated Dye 2 for each neuron (Fig. 3E-H). The mean fractions of potentiated spines per neuron were not significantly different between the CFC and context-only groups, but both groups were significantly higher than in the home cage control group (CFC: 1.6 ± 0.27%, mean ± s.e.m., n = 75,200 spines, 19 cells, 4 mice; context-only: 0.97 ± 0.12%, mean ± s.e.m, n = 74,801 spines, 17 cells, 4 mice; home cage control: 0.36 ± 0.06%, mean ± s.e.m, n = 43,659 spines, 12 cells, 3 mice; Fig. 3I). In the CFC mice, a subset of neurons had an elevated fraction of potentiated spines compared to context-only controls (5 of 19 neurons in CFC mice had a higher fraction of potentiated spines than all 17 neurons from context-only controls, Fig. 3I). Furthermore, the Dye 2 fluorescence in potentiated spines in the CFC group was significantly brighter than in either the home cage control or the context-only group (CFC: 1,000 ± 35 counts, mean ± s.e.m., n = 1016 spines from 19 cells, 4 mice; context-only: 830 ± 38 counts, mean ± s.e.m., n = 683 spines from 16 cells, 4 mice; home cage control: 789 ± 36 counts, mean ± s.e.m., n = 215 spines from 12 cells, 3 mice; Fig. 3J). Finally, within the CFC group, the level of Dye 2 in potentiated spines correlated with the fraction of potentiated spines on a cell-by-cell basis (*R* = 0.48, *P* = 0.04; Fig. S10A). These results imply that the conditioned stimulus elevated the percentage of potentiated spines and the degree of AMPAR exocytosis in a subset of neurons.

We next compared AMPAR exocytosis and cFos expression at the level of individual cells (Fig. 3K-O and Fig. S10B). Mice exposed to CFC had more cFos expression compared to either context-only or home cage controls (CFC: cFos level 1.1×10^4^ ± 1.1×10^2^ A.U., mean ± s.e.m., n = 7,208 cells, 4 mice; context-only: 8.0×10^3^ ± 1.3×10^2^ A.U., mean ± s.e.m, n = 5,074 cells, 4 mice; home cage control: 5.8×10^3^ ± 1.3×10^2^ A.U., mean ± s.e.m, n = 1,922 cells, 3 mice; Fig. 3M, Fig. S10), consistent with previous reports ^47^. We plotted the fraction of spines with high AMPAR exocytosis as a function of the cFos expression, cell by cell. These two quantities were highly correlated within the CFC group (*R* = 0.90, *P* < 0.0001), but not in the context-only group, nor in the home cage control group (Fig. 3N, O). To rule out imaging artifacts, we verified that the fraction of potentiated spines, the cFos intensity, and the total number of detected spines were independent of the imaging depth below the slice surface in either the CFC or context-only groups (Fig. S11). Moreover, the total number of spines (labeled with Dye 1) did not correlate with cFos intensity (Fig. S12). Together, these results establish that during the formation of associative memory, CA1 neurons with higher activity (as reported by cFos) exhibited higher AMPAR exocytosis compared to low-cFos neurons from the same animal.

### Mapping sub-cellular patterns of plasticity

We then analyzed the sub-cellular distributions of spines with high AMPAR exocytosis. We segmented the dendrites and registered all spines to their corresponding dendrites (Fig. S13). AMPAR exocytosis was more prevalent in perisomatic than in distal spines (Fig. 4A and Figs. S14-16), with a bias toward basal over apical dendrites (Fig. 4B and Fig. S17). We did not observe any difference in the spatial distribution of AMPAR exocytosis between CFC vs control mice, nor between high and low cFos expressing cells within each experimental group (Fig. S18).

**Fig. 4.**
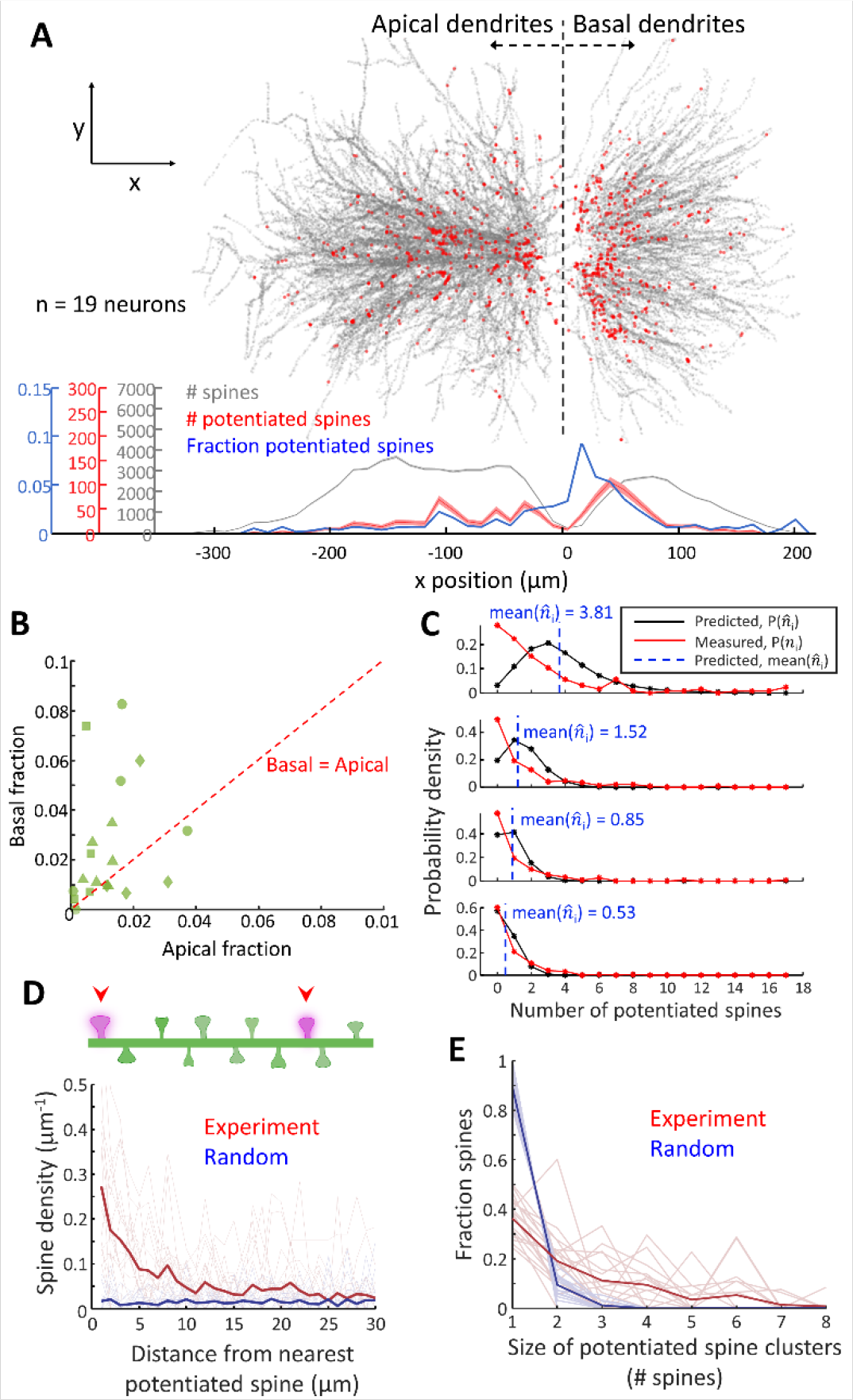
Sub-cellular distribution of AMPAR exocytosis in CA1 pyramidal neurons. **(A)** Distribution of spines with high HT-GluA1 exocytosis as a function of distance from the *stratum pyramidale*. (Top) Overlapped images of all identified spines (grey) and spines with high HT-GluA1 exocytosis (red) from the CFC group. (Bottom) Density profiles of all spines (grey), potentiated spines (red), and fraction of potentiated spines (blue). Shading represents count ± sqrt(count). **(B)** Fraction of potentiated spines in basal vs. apical dendrites in neurons from CFC group (*n* = 19 neurons). **(C)** Distributions of the measured (*n*_i_) and predicted 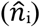 number of potentiated spines in individual dendrite segments from the CFC group (*n* = 1476 dendrite segments, 19 neurons). 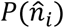 was evaluated from 500 random allocations of potentiated spines among dendrites of a given branch order within each cell. The plots show distributions for groups of branches with similar 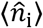 (regardless of branch order) and sorted by decreasing mean 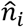 from top to bottom. **(D)** Density profile of potentiated spines as a function of distance from the nearest potentiated spine. The single-cell profiles are plotted with light colors. Random: simulation where the same number of potentiated spines were distributed randomly and independently among all detected spines (*n* = 19 neurons from the CFC group). **(E)** Fraction of potentiated spines clusters of different sizes from the CFC group (*n* = 19 neurons). The single-cell profiles are plotted with light colors. Random defined as in **(D)**.

We next sought to assess whether individual dendritic branches had either an excess or deficit in number of potentiated spines. To remove the overall dependence of plasticity on distance from the soma, we compared dendrites of equal branch order. Within each neuron and each branch order, we quantified the number of potentiated spines, *n*_i_, and the total number of spines, α_i_, in each dendritic segment *i*. We also calculated the total number of potentiated spines, *N*, and all spines, *A*, at the corresponding branch order. If potentiated spines were allocated randomly, one would expect *n*_i_/*N* ≈ α_i_/*A*. We performed a stochastic simulation to estimate the distribution of expected 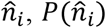, under the random allocation hypothesis for each branch, and compared to our data. Compared to the simulated random allocation, the data contained an excess of branches with zero potentiated spines, and an excess of branches with more-than-expected potentiated spines (Fig. 4C). These findings point to the presence of “silent” branches (with respect to plasticity), as well as a dendrite-level clustering of plasticity.

Finally, we examined whether potentiated spines showed fine-scale clustering within individual dendritic branches. For each neuron, we calculated the inter-spine contour distance between all pairs of spines. The pairwise distance distribution between potentiated spines showed a decay with a length constant of 8.0 μm (95% confidence interval [6.5 10]). In a simulation where we took the same number of potentiated spines and distributed them randomly and independently among all spines, the corresponding distribution for randomly selected spines was flat (Fig. 4D, Fig. S19A). We then quantified the distribution of cluster size, i.e. the number of potentiated spines within each cluster (Fig. 4E, Fig. S19B, materials and methods). In CFC-treated mice, potentiated spines were more likely to be in clusters of ≥ 2 spines compared to the simulated random allocation (potentiated: 57 ± 4.3% of spines were in clusters, mean ± s.e.m.; randomly allocated: 7.2 ± 1.5% clustered, mean ± s.e.m.; *P* = 2.0×10^−4^ by Wilcoxon signed-rank test). Clustering of potentiated spines was similar in context-only control mice (context-only: 54 ± 3.7%, mean ± s.e.m.; *P* = 0.19 by two-sided Wilcoxon rank-sum test, Fig. 4E and Fig. S20B). These analyses illustrate how pulse-chase HT labeling can map sub-cellular details of synaptic potentiation.

## Discussion

A longstanding question in the engram field has been to identify the biophysical mechanisms by which cFos-positive engram cells modulate circuit dynamics to encode a memory ^30^. Our work shows that cFos expression can serve as a surrogate for synaptic plasticity, and thereby connects the seemingly disparate bodies of work on engram cells and on synaptic encoding of memories.

Mice exposed to the same context without aversive stimulus exhibited less AMPAR exocytosis, lower cFos expression, and did not exhibit a correlation between AMPAR exocytosis and cFos. These results suggest an important role for reinforcement signals in mediating both AMPAR exocytosis and cFos expression. Our experiments do not distinguish the relative roles of intrinsic excitability ^48–50^ vs. variations in synaptic inputs ^51,52^ as the primary trigger for selecting the specific engram cells.

Dendrites of CA1 pyramidal neurons receive synaptic inputs from multiple pathways arranged in a laminar distribution. It has been proposed that distal input onto apical dendrites from entorhinal cortex act as an instructive signal, which modulates plasticity that primarily occurs at the proximal inputs from CA3 ^53^. Our results are consistent with this picture and further show that under our experimental paradigms there was little potentiation of the distal inputs. Choi and coworkers showed that in CFC-treated mice, synapses between CA3 engram cells and CA1 engram cells were enlarged ^23^. We observed that in CA1, potentiated spines were preferentially located in proximal dendrites, with a bias for basal vs. apical, which matches the distribution of CA3 inputs ^54^. Our results are thus consistent with the observations of Choi and coworkers, and further show that an increase in AMPAR density occurred in the 90 minutes after the CFC treatment. Attenuation of backpropagating action potentials, which can activate NMDA receptors, may also have contributed to preferential plasticity near the soma ^55^. Our results are also consistent with prior findings that synaptic plasticity in cortical pyramidal neurons is differentially regulated between lamina ^56^.

We observed clustering of synaptic plasticity at the branch level, aligning with previous theoretical predictions and *in vivo* observations ^57^. This finding supports the possibility that individual dendrites can serve as units for memory allocation. This dendrite-level plasticity may arise from compartmentalized calcium influx, which can initiate post-synaptic signaling pathways associated with LTP ^58^. Finally we observed short-range (< 10 μm) clustering of potentiated spines, consistent with prior observations in cultured neurons and in superficial cortex *in vivo* ^10,59–61^. This clustering may originate from either co-activated synapses ^62^ or from diffusion of small GTPases between nearby spines ^63–66^. Clustered plasticity has been proposed to facilitate local dendritic spike generation ^31,67^.

Pulse-chase labeling with membrane-permeable HaloTag ligand dyes was recently used to map protein turnover in the brain ^24^. Labeling of surface-exposed proteins with impermeable dyes probes membrane trafficking, which is an important post-translational means of regulating activity. We expect that HT-GluA1 can be useful for studies on memory, and the EPSILON approach can be adapted for other transmembrane proteins too, using either viral, transgenic^15^ or CRISPR-based knock-in^68–70^ approaches.

## Acknowledgments

We thank Andrew Preecha and Shahinoor Begum for technical assistance. We thank Richard Huganir for sharing plasmids. We thank Michael Tadross, Ryohei Yasuda, and Bernardo Sabatini for helpful discussions. This work was supported by a grant from Schmidt Futures and NIH grant 1-R21-EY033669. J.D.W.-C. is a Merck Awardee of the Life Sciences Research Foundation. J.B.G, and L.L. are supported by the Howard Hughes Medical Institute.

## Materials and Methods

### Animals

All experiments were performed on 8-12 week-old male and female C57BL/6 and CD-1 mice purchased from Charles River Laboratories. All animal procedures were in accordance with the US National Institutes of Health *Guide for the Care and Use of Laboratory Animals* and were approved by the Institutional Animal Care and Use Committee at Harvard University.

### DNA constructs

Standard methods of molecular cloning were used to create the constructs. The myc-GluA2 plasmid was provided by the Richard Huganir lab at Johns Hopkins University ^10,12,13^. Plasmids and sequences created for this project are on Addgene.

We chose to overexpress HT-GluA1, rather than knocking HT into the endogenous locus to mirror previous results with the analogous SEP-GluA1 construct. Overexpression of SEP-GluA1 in both cultured neurons and *in vivo* did not perturb the neuron’s intrinsic or synaptic properties ^10,12^ while SEP-GluA1 knock-in decreased GluA1 mRNA and protein expression ^15^.

#### pTL024 (Addgene 192517)

Signal sequence-HaloTag-GluA1 (abbreviated HT-GluA1), driven by human synapsin 1 gene (hSynI) promoter.

Plasmid pTL024 was assembled from three fragments: the N terminal signal sequence, the HaloTag protein, and GluA1. The N terminal signal sequence was amplified from pCI-SEP-GluR1, Addgene 24000. The HaloTag protein was amplified from Voltron, Addgene 119033. GluA1, along with a 15 amino acid linker, was amplified from CAG::SEP-GluA1, a gift of Prof. Richard Huganir ^10,12,13^. The fragments were assembled together with Gibson Assembly ^71^ and cloned into pLenti hSynI vector (HT075 from ^72^).

### Synthesis of Alexa Fluor 647-HaloTag ligand (AF647-HTL)

Alexa Fluor 647 NHS ester (5 mg, 4.0 μmol) and HaloTag amine (O2) ligand (1.3 mg, 6.0 μmol, 1.5 eq) were combined in DMF (1 mL), and DIEA (3.5 μL, 20.0 μmol, 5 eq) was added. The reaction was stirred at room temperature for 18 h while shielded from light. It was subsequently purified by reverse phase HPLC (5–30% MeCN/H_2_O, linear gradient, with constant 0.1% v/v TFA additive; 20 min run; 42 mL/min flow; Gemini-NX 5 μm, 30 × 150 mm) to provide 3.2 mg (68%) of the title compound as a blue solid. Analytical HPLC: t_R_ = 13.9 min, >99% purity (10–50% MeCN/H_2_O, linear gradient, with constant 0.1% v/v TFA additive; 20 min run; 1 mL/min flow; Eclipse XDB 5 μm, 4.6 × 150 mm; detection at 650 nm; ESI, positive ion mode); MS (ESI) calculated for C_46_H_67_ClN_3_O_15_S_4_ [M]^+^ 1064.3, found 1064.1; MS (ESI) calculated for C_46_H_68_ClN_3_O_15_S_4_ [M+H]^2+^ 532.7, found 532.6.

### Primary neuron culture

14-mm glass-bottom dishes were incubated with 40 μg ml^-1^ of poly-D-lysine (PDL) in phosphate-buffered saline (PBS) for 1 h at room temperature, followed by an overnight incubation at 4 °C with 20 μg ml^-1^ laminin (Thermo Fisher Scientific, 23-017-015). After being thoroughly washed with PBS, hippocampi were dissected from embryonic day 18 (E18) rats and resuspended in BrainPhys medium (BPNM, STEMCELL Technologies, 05790) supplemented with 2% SM1 (STEMCELL Technologies, 05792), 5 mM L-glutamine (STEMCELL Technologies, 07100), and 35 μg ml^-1^ of L-glutamic acid (Sigma-Aldrich, 49449) to a final concentration of 3.0 x 10^6^ cells per milliliter. The neurons were then plated at a density of 30,000 cells per cm^2^ on the pre-treated glass-bottom dishes, followed by the addition of 2 ml of BPNM with 2% SM1 (BPNM/SM1). The health of the neurons was monitored daily from days *in vitro* 1 (DIV1) to DIV7, and the medium in each dish was replaced with 37 °C fresh BPNM/SM1 medium every 3-4 days.

### Virus packaging

In-house preparations of lentivirus were used in this study, based on a second-generation lentivirus packaging system. The detailed methods of lentivirus production and packing were previously described ^73^.

### HT-GluA1 expression in cultured neurons

The hSyn::HT-GluA1 vectors were introduced to the neurons via lentiviral transduction at DIV7. The lentiviral vectors were added directly to the neuronal cultures in fresh BPNM/SM1 medium. The neuronal cultures were then incubated with the lentivirus-containing medium for 12 hours at 37 °C and 5% CO_2_, followed by a medium replacement with lentivirus-free medium. Experiments were conducted at DIV14.

### Manders’ overlap coefficients

The Manders’ coefficients (M1 and M2) were calculated between two-channels (HT-GluA1 and PSD-95) for each neuron as previously described ^74^ using MATLAB. Briefly, the images in each color channel were binarized with a threshold set by the Otsu algorithm. M1 and M2 were calculated via:

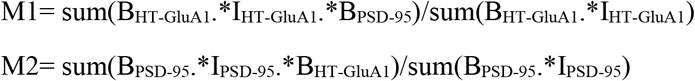

where I denotes the original image, B denotes a binary mask obtained with a threshold set by the Otsu algorithm. ‘.*’ is elementwise multiplication.

### *In vitro* labeling kinetics

Cultured neurons expressing HT-GluA1 at DIV14 were incubated with 100 nM AF_488_-HTL or 1μM JF_549i_-HTL in fresh BPNM/SM1 medium for different time durations (5 seconds, 30 seconds, 5 minutes, and 30 minutes for 100 nM AF_488_-HTL; 5 seconds, 10 seconds, 30 seconds, 2 minutes, 5 minutes, and 10 minutes for 1μM JF_549i_-HTL) at 37°C and 5% CO_2_. The neurons were then washed twice with PBS for 1 minute each and fixed with 4% paraformaldehyde. The fixed neuron cultures were then washed in PBS for 24 hours. All media were heated to 37 °C before use.

### *In vitro* turnover rate

Cultured neurons expressing HT-GluA1 at DIV14 were incubated with 100 nM AF_488_-HTL in fresh BPNM/SM1 medium for 5 minutes at 37 °C and 5% CO_2_. The neurons were then washed twice with PBS for 1 minute each and the medium was replaced with dye-free medium. To label new surface HT-GluA1, the medium was replaced with 1 μM JF_549i_-HTL in fresh BPNM/SM1 medium at different time points. After incubating for 5 minutes at 37 °C and 5% CO_2_, neurons were washed twice with PBS for 1 minute each and fixed with 4% paraformaldehyde. The fixed neuron cultures were then washed in PBS for 24 hours. All media were heated to 37 °C before use.

### Pulse-chase labeling in cultured neuron upon chemical LTP

Cultured neurons expressing HT-GluA1 at DIV14 were exposed to 100 nM AF_488_-HTL in Mg^2+^-free ACSF at 37 °C and 5% CO_2_ for 5 minutes. Afterward, the neurons were washed twice with Mg^2+^-free ACSF for 1 minute each and treated with Mg^2+^-free ACSF containing 100 nM rolipram, 50 μM forskolin, and 100 μM picrotoxin for 20 minutes at 37 °C and 5% CO_2_. To label new surface HT-GluA1 upon cLTP, the neurons were then incubated with 1 μM JF549i-HTL in Mg^2+^-free ACSF for 5 minutes at 37 °C and 5% CO2 before being washed twice with Mg^2+^-free ACSF for 1 minute each and fixed with 4% paraformaldehyde. The fixed neuron cultures were washed in PBS for 24 hours. The negative control cultures were exposed to Mg^2+^-free ACSF without rolipram, forskolin, and picrotoxin. All media were pre-incubated with 5% CO_2_ for 30 minutes and heated to 37 °C before use.

### *In utero* electroporation

Progenitor cells in layer 2/3 of the embryonic mouse brain were transfected using *in utero* electroporation. Pregnant CD-1 mice were used and DNA solution containing Fast Green was injected into the lateral ventricle of each embryo through a pulled-glass pipette. Electric pulses were applied using 5-mm Pt electrodes. The angle of electrodes was adjusted to target the specified brain region (barrel cortex or hippocampal CA1). The electroporation protocol comprised five pulses of 35 V, with a frequency of 1 Hz and a duration of 50 ms. The DNA solution used contained HT-GluA1, myc-GluA2 at a ratio of 1:1 (final concentration: 2 μg/μL each). For the co-expression with GPI-GFP, the DNA solution contained HT-GluA1, myc-GluA2, and GPI-GFP at a ratio of 2:2:1 (final concentration: 2 μg/μL for HT-GluA1 and myc-GluA2; 1 μg/μL for GPI-GFP).

### Patch-clamp electrophysiology

Coronal brain slices (300 μm) were prepared from CD-1 mice of either sex between postnatal days P14 and P16. IUE was used for HT-GluA1 expression in cortical layer 2/3 neurons. Standard whole-cell recording was performed at 34 °C during a continuous perfusion at 2 ml min^−1^. The perfusion buffer contained 124 mM NaCl, 3 mM KCl, 26 mM NaHCO_3_, 1.25 mM NaH_2_PO_4_, 2 mM MgCl_2_, 15 mM D-glucose, and 2 mM CaCl_2_ (saturated with 95% O_2_ and 5% CO_2_). Cortical layer 2/3 neurons were visualized using a custom-built upright microscope. The whole-cell internal solution contained 8 mM NaCl, 130 mM KMeSO_3_, 10 mM HEPES, 5 mM KCl, 0.5 mM EGTA, 4 mM Mg-ATP, 0.3 mM Na_3_-GTP. The pH was adjusted to 7.2–7.3 with KOH and osmolarity was set to 290–295 mOsm l^−1^. Borosilicate glass pipettes were used with a resistance of 3-5 MΩ (1B150F-4; WPI). Patch clamp recordings were acquired and filtered at 10 kHz with the internal Bessel filter using a Multiclamp 700B (Molecular Devices) and digitized with PCIe-6323 (National Instruments) at 100 kHz. Following the whole-cell configuration, membrane capacitance (C_m_), and membrane resistance (R_m_) were estimated under voltage clamp mode. Resting membrane potential, rheobase, and spike rates were measured under current clamp mode. Rheobase was defined as the minimum current step (in 500 ms duration) required to trigger at least one spike. Whole-cell recordings were monitored and analyzed in MATLAB.

To measure evoked EPSCs, voltage-clamp experiments were performed with a stimulating electrode (CBAPB50, FHC) placed 100-200 μm laterally to activate layer 2/3 inputs. Holding potential was -70 mV. The whole-cell internal solution contained (in mM) 8 NaCl, 130 CsMeSO_3_, 10 HEPES, 0.5 EGTA, 4 Mg-ATP, 0.3 Na_3_-GTP, 5 QX-314, 0.1 spermine. The pH was adjusted to 7.2-7.3 with CsOH and osmolarity was set to 290-295 mOsm l^−1^. The perfusion buffer contained 50 μM picrotoxin (Thermo Scientific) and 10 μM (+)-bicuculline (Enzo Life Sciences) to prevent GABA_A_ receptor-mediated transmission. After a stable baseline of at least 5 min, the input-output relationships were estimated by varying the stimulus intensity from 3 to 15 V in 3 V increments (0.1 ms duration). Stimulation frequency was 0.1 Hz. For measuring the AMPAR/NMDAR ratio, cells were clamped at a holding potential of - 70 mV to measure the peak of AMPAR-mediated synaptic transmission. NMDAR-mediated currents were estimated at 75 ms after the stimulus onset at a holding potential of +40 mV. Example traces were an average of 5 consecutive responses, collected from typical experiments (stimulus artifacts were blanked for clarity). Experiments were accepted for analysis if series resistance values were < 20 MΩ and varied by <15% throughout the experiment.

### Installation of dye injection port

Male or female CD-1 mice expressing HT-GluA1 via *in utero* electroporation were selected at age 9-11 weeks and anesthetized using a ketamine/dexmedetomidine solution. To maintain body temperature at 36-37 °C, a heating pad (WPI, ATC2000) was placed under the mouse, and ophthalmic eye ointment was applied to the mouse’s eyes to keep them moist. Surgical coordinates were identified as indicated in Figure S6, and a hollow titanium ring with an outer diameter of 5 mm, an inner diameter of 2.1 mm, and a thickness of 0.35 mm was attached to the skull surrounding the target coordinates with dental cement (C&B metabond, Parkell, No. 242-3200). Holes were drilled through the skull, and the titanium ring was filled with silicone gel which was removed before dye injection and refilled after injection. The mice were returned to their home cages after recovering from anesthesia.

### *In vivo* turnover rate measurement

One week after installation of the injection port at the barrel cortex, mice (age 10-12 weeks) were anesthetized with isoflurane and placed on a surgical platform. At every indicated injection point in Figure S6, 80 nL of AF_647_-HTL (1 μM) was injected at a rate of 100 nL/min. Then, 80 nL of JF_549i_-HTL (10 μM) was injected at the same points 20 minutes after the AF_647_-HTL injection, also at the same rate. After 20 minutes from the end of the dye injection, the mice were returned to their home cages. Later, at different timepoints, the mice underwent the second JF_549i_-HTL injection. At the same injection points as previously described, 80 nL of JF_549i_-HTL (10 μM) was injected at the same rate. After 20 minutes from the end of the dye injection, the mice were euthanized and prepared for brain slicing.

### Acute whisker stimulation with pulse-chase labeling

Whisker stimulation sessions were conducted one week after injection port installation. On the day of stimulation, 2.7 mg/kg of chlorprothixene hydrochloride dissolved in PBS was intraperitoneally injected before anesthetic induction. Mice (10-12 weeks) were anesthetized with isoflurane on the surgical platform and injected at each point shown in Figure S6 with JF_549i_-HTL (80 nL, 1 μM) at a rate of 100 nL/min. After 20 minutes, AF_488_-HTL (80 nL, 10 μM) was injected at the same points. After another 20 minutes from the end of dye injection, the contralateral whiskers were trimmed in a chessboard pattern ^39–41^. Spared whiskers were deflected at 10 Hz for 180 minutes with a rotary whisker stimulator. Mice in the control group were placed under the same experimental conditions, but all contralateral whiskers were trimmed. During whisker stimulation, isoflurane level was kept at ∼0.5% to maintain shallow anesthesia. After whisker stimulation, the mice underwent the second injection of AF_488_-HTL (80 nL, 10 μM) as previously described. After 20 minutes from the end of the dye injection, the mice were euthanized and prepared for brain slicing.

### Slice preparation for confocal imaging

Mice were overdosed with isoflurane until their breathing ceased. Their brains were then promptly extracted. The brains were sliced into 300 μm coronal sections using a vibratome (Leica, VT1200S) and then fixed in 4% paraformaldehyde for 24 hours at 4 °C. The fixed slices were additionally washed in PBS for 48 hours on a shaker at room temperature. The slices were mounted (VECTASHIELD PLUS Antifade Mounting Medium, Vectorlabs, H-1900-10) to be imaged on confocal imaging systems.

### Contextual fear conditioning with pulse-chase labeling

In the five days prior to CFC treatment, each mouse was housed alone and habituated to the investigator and anesthesia chamber without isoflurane. On the day of conditioning, the mouse (age 10-12 weeks) was anesthetized for 3 hours with 1.0% isoflurane on the surgical platform and injected with AF_647_-HTL (80 nL, 1 μM) at a rate of 100 nL/min at every injection point shown in Figure S6. After 20 min, JF_549i_-HTL (80 nL, 10 μM) was injected at the same points. The mouse was then returned to its home cage to recover from anesthesia. After 20 minutes, the mouse underwent conditioning sessions that lasted 300 s. For the first 150 s, the mouse was allowed to explore the conditioning chamber freely. Starting at 150 s and repeating every 30 s for a total of five shocks, the mouse was given 0.7 mA foot shocks of 2-second duration. Mice in the context-only group were placed in the same chamber for 300 seconds without any shocks. Mice in the home cage control group remained in their home cage. After 90 minutes from the end of the conditioning session, the mouse was anesthetized with isoflurane on the surgical platform for the second JF_549i_-HTL injection. 80 nL of JF_549i_-HTL (10 μM) was injected into the injection points at the same points and rate as previously described. After 20 minutes from the end of the dye injection, the mouse was euthanized and prepared for brain slicing.

For the experiments in Figure S8, the same conditioning sessions (with and without shocks) were conducted, except the dye injection steps were skipped. 1 day after conditioning, mice were exposed to the same context and their freezing levels were measured via video analysis of the first 180 seconds from re-exposure. Animal motion was tracked using MATLAB code. Briefly, the center of mass of the mouse was tracked for every frame using the regionprops function. Then, the mouse’s speed was calculated between each frame, and the time duration during which the speed was slower than 0.03 m/s was counted as freezing time.

### cFos immunohistochemistry

After the brain was sliced, fixed, and washed as described above (slice thickness = 150 μm), the fixed slices were permeabilized with PBST (1% Triton-X in PBS) for 24 hours at room temperature on a shaker. The slices were then blocked with 5% bovine serum albumin (BSA) in PBST for 1 hour on a shaker. For cFos immunostaining, the slices were incubated with a rat anti-cFos primary antibody (1:1000 dilution in 1% BSA in PBST, Synaptic Systems, 226 017) for 24 hours at room temperature on a shaker. Slices were then washed in PBST for 20-min (x3), followed by 2-hour incubation with secondary antibody (1:500 in 1% BSA in PBST; AF_488_ anti-rat, Invitrogen, A-11006) at room temperature. After two additional 20-minute washes and one 24-hour wash in PBST at room temperature, the mounted slices were imaged on a confocal microscope.

### Confocal imaging of HT-GluA1 expressed in cultured rat hippocampal neuron

Confocal images of fixed neuron cultures were acquired using LSM 980 with 20× water immersion objective. AF_488_, JF_549i_, AF_647_ were excited with 488-nm, 561-nm, and 633-nm lasers, respectively. Pixel size was 0.052 μm by 0.052 μm and the size of each region of interest (ROI) was 424.27 μm by 424.27 μm (8192 pixels by 8192 pixels). Pixel time was 0.51 μs. The same imaging conditions were used throughout all experiments.

### Confocal imaging of HT-GluA1 expressed in brain slice

Confocal images of fixed brain slices were acquired using LSM 980 with 20× water immersion objective in Z-stack. Images in Figure 2A (right), 3A (right), and 3D were acquired with 63× oil immersion objective. DAPI, GFP, AF_488_, JF_549_, JF_549i_, and AF_647_ were excited with 405-nm, 488-nm, 488-nm, 561-nm, 561-nm, 633-nm lasers, respectively. Non-specific autofluorescent artifacts were excited with 488-nm for Figure 2 and 561-nm for Figure 3 and the detection ranges were from 500-nm to 553-nm for Figure 2F-I, from 651-nm to 695-nm for Figure 2J-N, and from 500-nm to 553-nm for Figure 3. Voxel sizes were 0.052 μm by 0.052 μm by 2 μm for Figure 2 (except for Figure 2A(right)) and 0.052 μm by 0.052 μm by 1 μm for Figure 3 (except for Figure 3A(right)). For Figure 2A(right) and 3A(right), the voxel size was 0.016 μm by 0.016 μm by 0.23 μm. Pixel time was 0.51 μs for all experiments. The same imaging conditions were used for all samples within each set of experiments.

### Image processing and data analysis

MATLAB, ImageJ, and Imaris were used for image processing. All images were denoised with 2D gaussian filter before analysis. The same kernel sizes were applied to all data within each set of experiments.

#### (1) Spine detection

The Imaris background subtraction function was applied to each image. The locations and fluorescence intensity values of spines were then determined by analyzing the intensity of the Dye 1 channel using the Imaris spot detection function (threshold: ‘quality’ above 130 for Figure 1, 600 for Figure 2, and 400 for Figure 3). For Figure 1, the dendrites were manually traced for each neuron, and spots located more than 5 μm away from the nearest dendrite were excluded from the analysis. For Figure 2 and 3, autofluorescent artifacts were identified by their intensities in the ‘autofluorescence detection channel’ (threshold: ‘quality’ above 300 for Figure 2F-I, 150 for Figure 2J-N and 1300 for Figure 3). Spots within 1 μm of the center of an autofluorescent artifact were excluded from analysis. To further reduce noise, spots whose average distance to 10 nearby spots was greater than 10 μm were filtered out. The same parameters were used throughout all experiments that required spine detection. For Figure 3 and 4, spines from different neurons in the same ROI were separated manually. Newly formed spines (i.e. synaptogenesis) were identified with the same criteria applied to the Dye 2 channel. Spots located within 4 μm from the nearest spine from Dye 1 were selected for the analysis. For background fluorescence analysis in Figure S7, 1 μm diameter spherical regions were randomly placed, with the constraints that they were at least 3 μm away from the nearest dendrite and from each other and at least 1 μm from the center of any autofluorescent puncta. The fluorescence intensities in both dye channels of these spherical regions were calculated following the same procedure as for spines.

#### (2) Threshold for defining potentiated spines

For each experimental condition, all spines from all ROIs were sorted into 300 bins based on their Dye 1 intensity. For each bin, we estimated the mean, μ, and standard deviation, σ, in the Dye 2 intensity of the non-potentiated spines from the bottom 95% of the Dye 2 distribution in that bin. A threshold was then set for each bin as μ + 5 σ, and spines with a Dye 2 intensity higher than the threshold were identified as ‘potentiated’. For Figure 1H and I, the thresholds were manually defined.

#### (3) cFos segmentation

Z-stack images were segmented based on the cFos channel (AF488) using Imaris and custom MATLAB code. The center coordinates of the nuclei were identified using Imaris (threshold: ‘quality’ above 1200), and a 20 μm by 20 μm image subset was defined around each center coordinate on the x-y plane. A binary mask was obtained from each image subset using the Otsu algorithm. Each binary mask underwent Euclidean distance transform followed by an extended minima transform. MATLAB function imextendedmin was used to filter out small local minima, and the remaining minima were imposed on the original binary mask. The resulting binary masks were segmented using the Watershed algorithm. Any masked area smaller than 45 μm^2^ was excluded from analysis. Background autofluorescence was calculated as the mean intensity of each non-masked area from the first Otsu thresholding. Background was subtracted from the mean intensity value within each masked area.

#### (4) Dendrite classification by branch order and spine assignment to dendrites

In Imaris, the coordinates of each dendrite were manually identified and extracted from each Z-stack image. For all dendrites, their branch orders were manually identified, and they were numbered according to their relative positions. For instance, #1340000 represents the fourth dendrite from the third second-ordered branch of the first first-ordered branch and its branch order is 3. The first digit of basal dendrite numbers ranged from 1 to 7, while the first digit of apical dendrite numbers ranged from 8 to 9. Custom MATLAB code was used to smooth these coordinates with the smoothdata function. Next, each dendrite was converted into 1000 equally spaced points using linear interpolation. The x, y, and z coordinates were imported from Imaris to MATLAB. For each spine, the dendrite closest to the spine was identified by determining the dendritic point with the shortest Euclidean distance to the spine. This dendrite was registered as the spine’s dendrite.

#### (5) Distribution of spines as a function of projected and contour distances from soma

The coordinates of all spines were transformed to match the dorsal-ventral (DV) axis of the neuron they belonged to. The centers of rotation for each neuron were manually determined by referring to the original confocal images and defined as the origin. In Figure 4A and S14, each spine point was orthogonally projected onto the corresponding DV axis crossing at the origin. The distance from the projected point to the origin was calculated and displayed. The distance values were binned into 50 groups, and the total number of spines, number of spines above threshold, and the fraction of spines above threshold were calculated for each bin. In Figure S15, the contour distance from the soma to each spine was calculated by adding the distance from the spine to one of the dendritic endpoints closer to the soma and the lengths of all dendrites between the endpoint and the soma.

#### (6) Calculation of pairwise distance distribution between potentiated spines

For each potentiated spine, the distances to all other potentiated spines within 30 μm dendrite contour distance were tabulated. The histogram of these values gave the pairwise distance distribution. For the random allocation, the same number of spines on each dendrite were randomly and independently selected and assigned as “pseudo potentiated spines”. The fraction of “pseudo potentiated spines” was calculated using the same method as above.

#### (7) Distribution of the expected number of potentiated spines per dendrite branch

Within each neuron and each branch order, we quantified the number of potentiated spines, *n*_i_, and the total number of spines, α_i_, in each dendritic segment *i*, and the total number of potentiated spines, *N*, and all spines, *A*, across all dendrite segments at the corresponding branch order. Next, we performed 500 simulations where we randomly distributed the *N* potentiated spines across the dendritic segments, with probability α_i_,/*A* of landing in segment *i*. We calculated the theoretical distribution of number of potentiated spines for each dendritic segment, 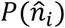, under the assumption of random and independent allocation within each branch order.

We then compiled all dendrite segments from all branch orders and all neurons and sorted them based on the expected mean number of potentiated spines derived from the simulations, 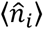. The sorted data were then divided into 10 groups of 130 dendrites each, and for each group, we computed the probability distribution of both the expected and measured number of potentiated spines. The four groups with the highest mean expected number of potentiated spines are presented in Figure 4C.

#### (8) Fraction of spine clusters with different sizes

We identified clusters of potentiated spines, first identifying clusters with the largest number of potentiated spines (*N* = 14 in our analysis) and then proceeding iteratively to analyze the remaining spines for clusters of size *N* – 1, *N* – 2, and so on. A cluster was defined as having *N* potentiated spines within a region containing 2*N* total spines on the dendrite. Once a cluster was identified and counted, its constituent spines were excluded from further counting to avoid duplication.

