## Supplementary Material for "Mapping memories: pulse-chase labeling reveals AMPA receptor dynamics during memory formation"

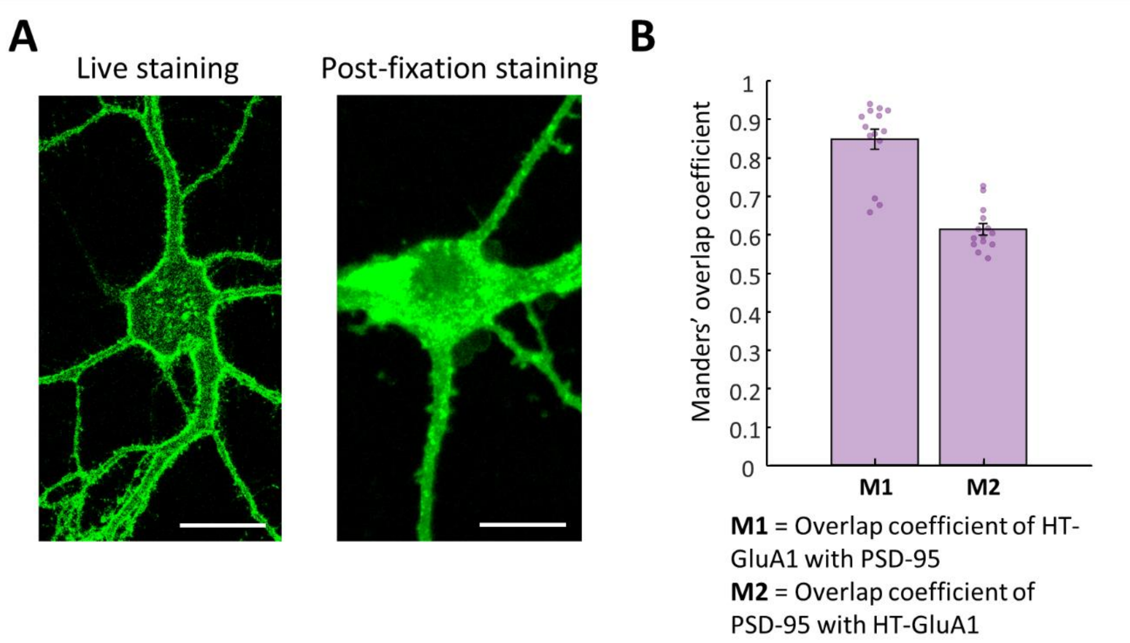


**Fig. S1. Trafficking of HT-GluA1 in cultured neurons.**

**(A)** Selective labeling of surface HT-GluA1 with membrane-impermeable HTL dye. Left: Cultured neuron expressing HT-GluA1 stained with membrane-impermeable $AF_{488}$-HTL while alive. Right: Another HT-GluA1 expressing cultured neuron stained with the same dye after fixation to permeabilize the cell membrane. Scale bars 10 μm. **(B)** Manders' overlap coefficient between HT-GluA1 and PSD-95 ($n$ = 14 cells from 3 cultures). Error bars show mean ± s.e.m.

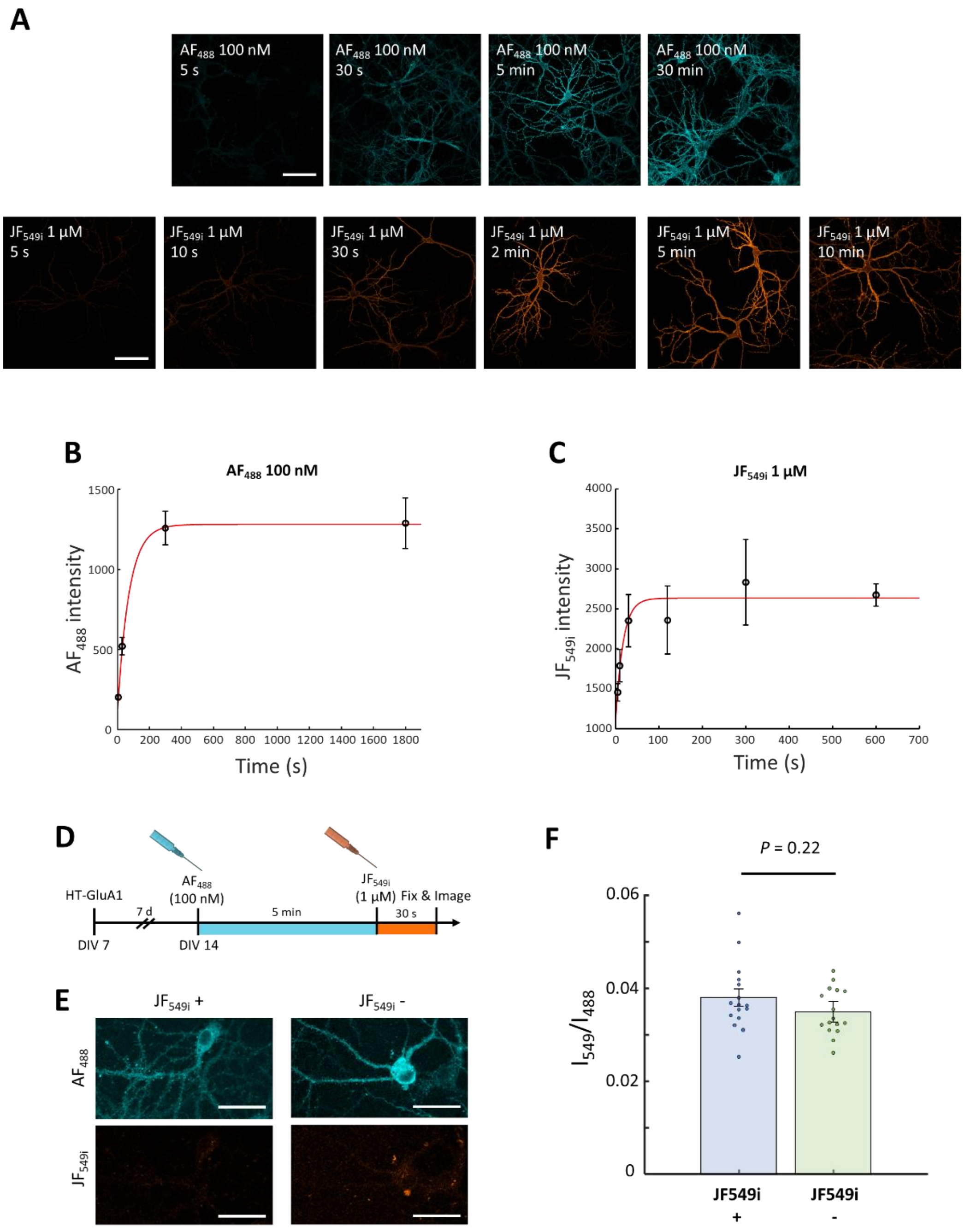


**Fig. S2. Labeling kinetics of membrane-impermeable HTL dyes on surface receptors.**

**(A)** Confocal images of fixed cultured neurons showing labeling of HT-GluA1 with $AF_{488}$-HTL (100 nM, cyan) or $JF_{549i}$-HTL (1 µM, orange) at different times after dye addition. Scale bars: 200 µm. **(B), (C)** Fluorescence vs. dye incubation time for cultured neurons expressing HT-GluA1 and treated with **(B)** $AF_{488}$-HTL (100 nM) or **(C)** $JF_{549i}$-HTL (1 µM). ($n$ = 5 cells for each timepoint). Data are represented as mean ± SD. Red: fitted curve. **(D)** Chase-dye labeling after saturating surface GluA1. Surface GluA1 was saturated by labeling with 100 nM $AF_{488}$-HTL for 5 minutes. 1 µM of $JF_{549i}$-HTL was immediately applied for 30 seconds. **(E)** Confocal images of fixed cultured neurons (left) with and (right) without $JF_{549i}$-HTL after saturation. Scale bars: 50 µm. **(F)** $JF_{549i}$-to-$AF_{488}$ intensity ratios ($n$ = 16 cells from 3 cultures for each experimental group). Error bars show mean ± s.e.m.

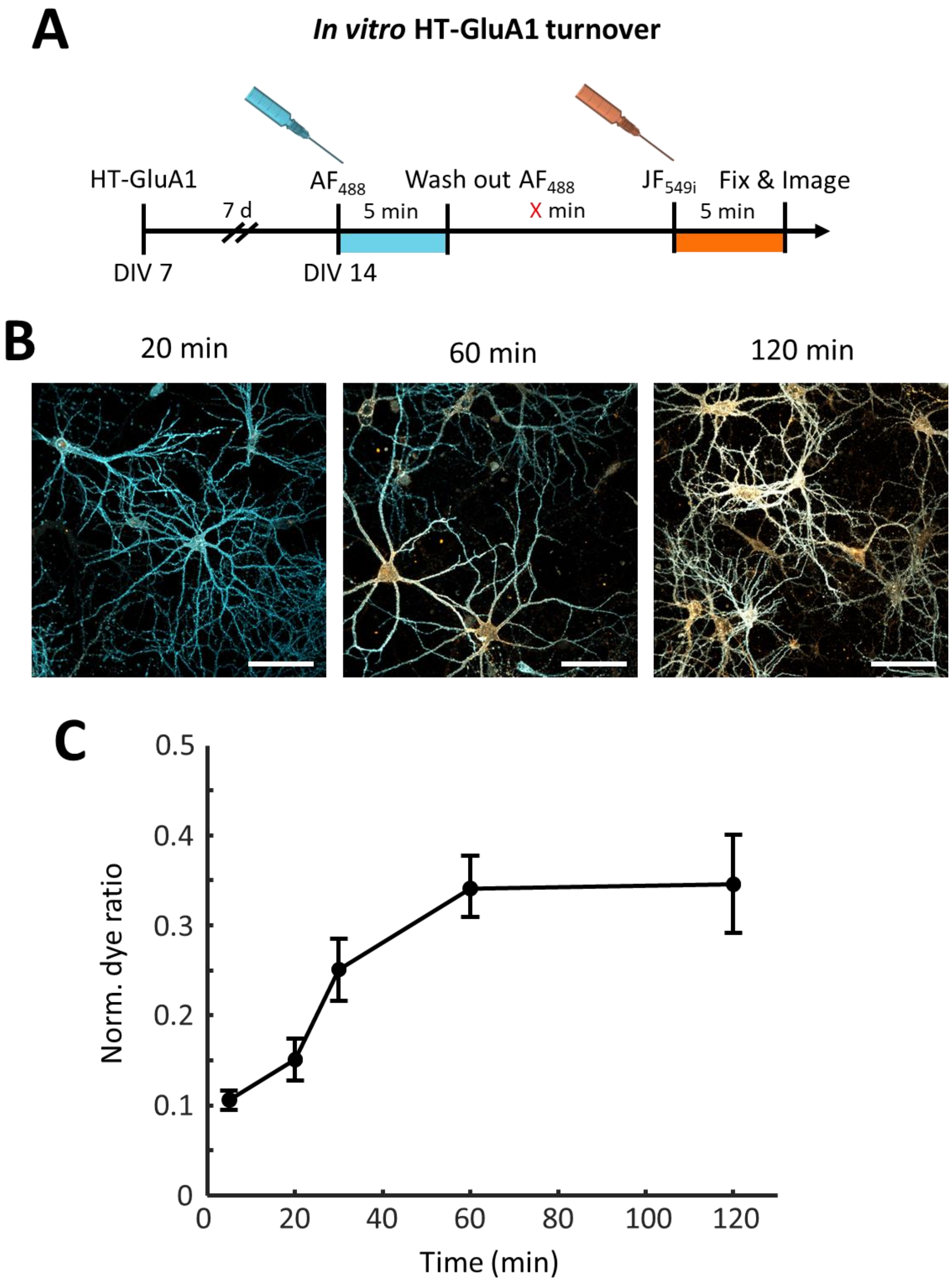


**Fig. S3. Pulse-chase measurement of HT-GluA1 turnover in cultured neurons.**

**(A)** Experimental timeline of surface HT-GluA1 turnover measurement with multi-color labeling. **(B)** Confocal images of fixed cultured neurons showing replacement of old ($AF_{488}$, cyan) with new ($JF_{549i}$, orange) AMPARs. Scale bars: 200 µm. Turnover occurred faster in soma and perisomatic neurites than in distal neurites. **(C)** Normalized dye intensity ratio at 5, 20, 30, 60, and 120-min (*n* = 5 cells for each timepoint). Normalized dye intensity ratio: $2^{nd}$ dye intensity / ($1^{st}$ dye intensity + $2^{nd}$ dye intensity). Data are represented as mean ± SD.

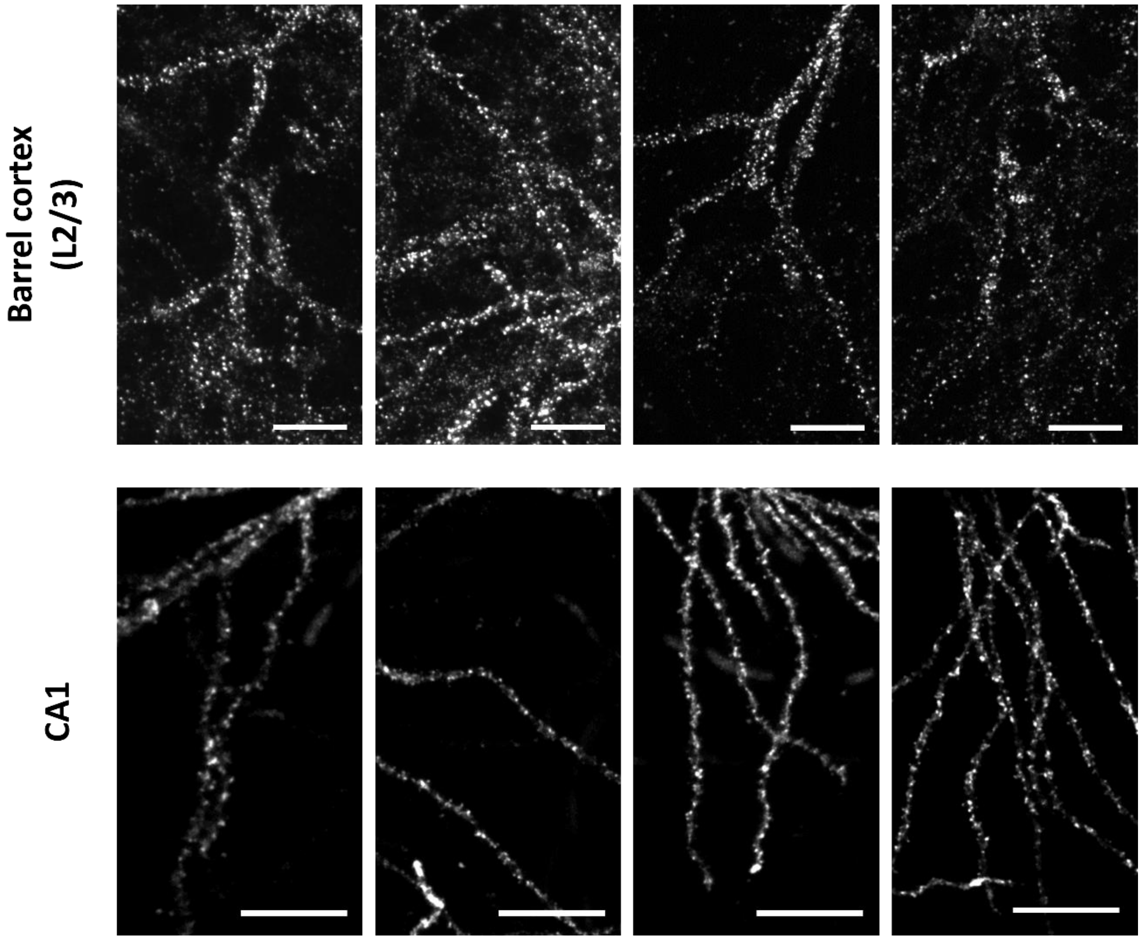


**Fig. S4. Confocal images of labeled neurons.**

Representative images of HT-GluA1-expressing pyramidal neurons stained with $AF_{647}$-HTL in mouse (top) barrel cortex layer 2/3 and (bottom) hippocampus CA1. Scale bars, 50 μm.

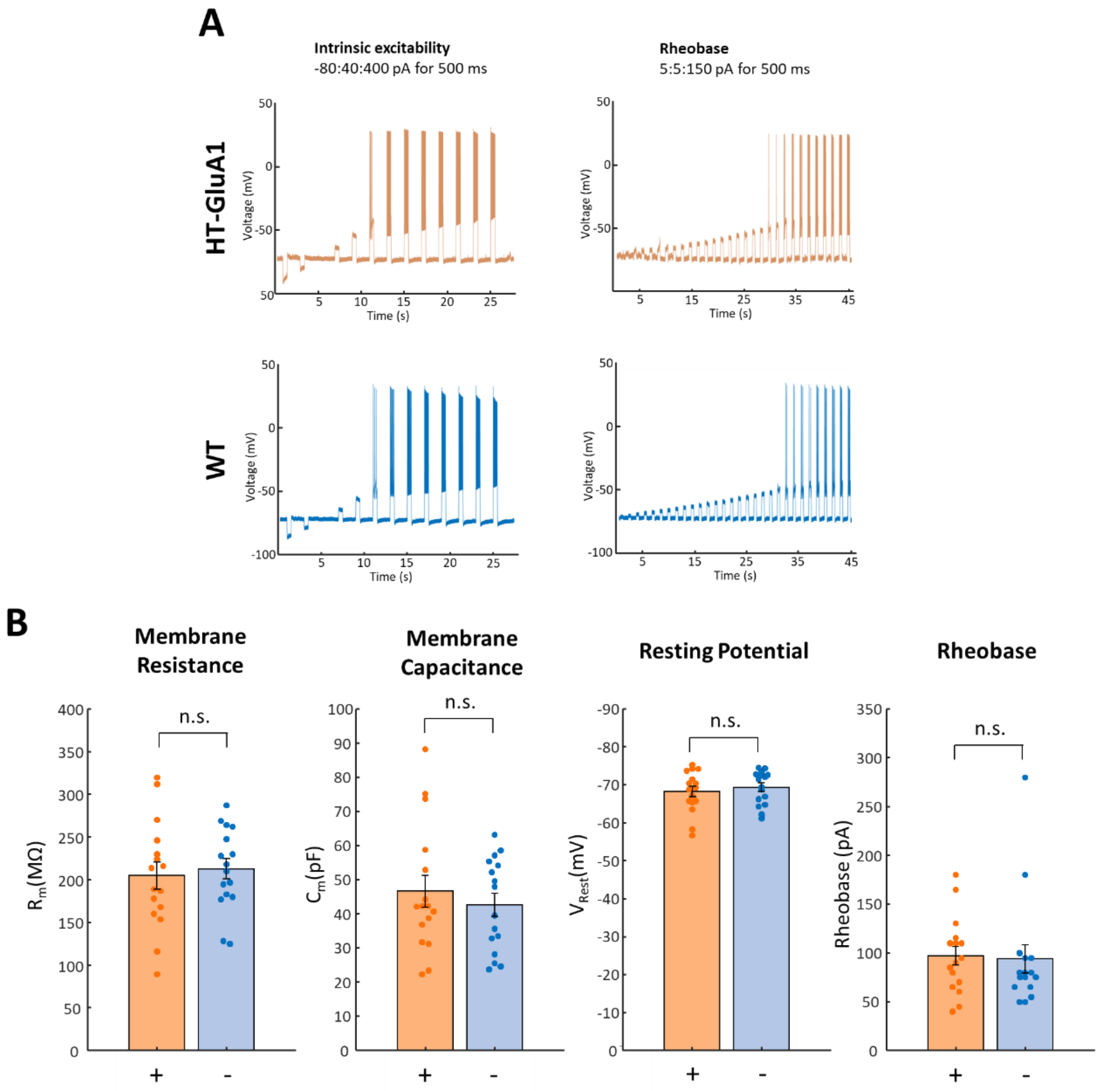


**Fig. S5. Patch-clamp characterization of HT-GluA1 expressing neurons in acute slice.**

HT-GluA1 expressing layer 2/3 pyramidal neurons in barrel cortex in acute brain slices were identified by staining with 1uM $JFX_{608}$-HTL. **(A)** Representative patch-clamp recordings in acute brain slice. **(B)** Measurements of electrophysiological properties of neurons with or without HT-GluA1 expression. Membrane resistance: 204 ± 16 MΩ vs 212 ± 12 MΩ, $P = 0.69$; membrane capacitance: 47 ± 5 pF vs. 43 ± 3 pF, $P = 0.50$; resting potential: −68.1 ± 1.4 mV vs. −69.2 ± 1.1 mV, $P = 0.53$; and rheobase 97 ± 9.8 pA vs. 94 ± 15 pA, $P = 0.86$ ($n$ = 13 neurons for each group). Error bars show mean ± s.e.m. n.s. not significant, two-sided Student's t-test.

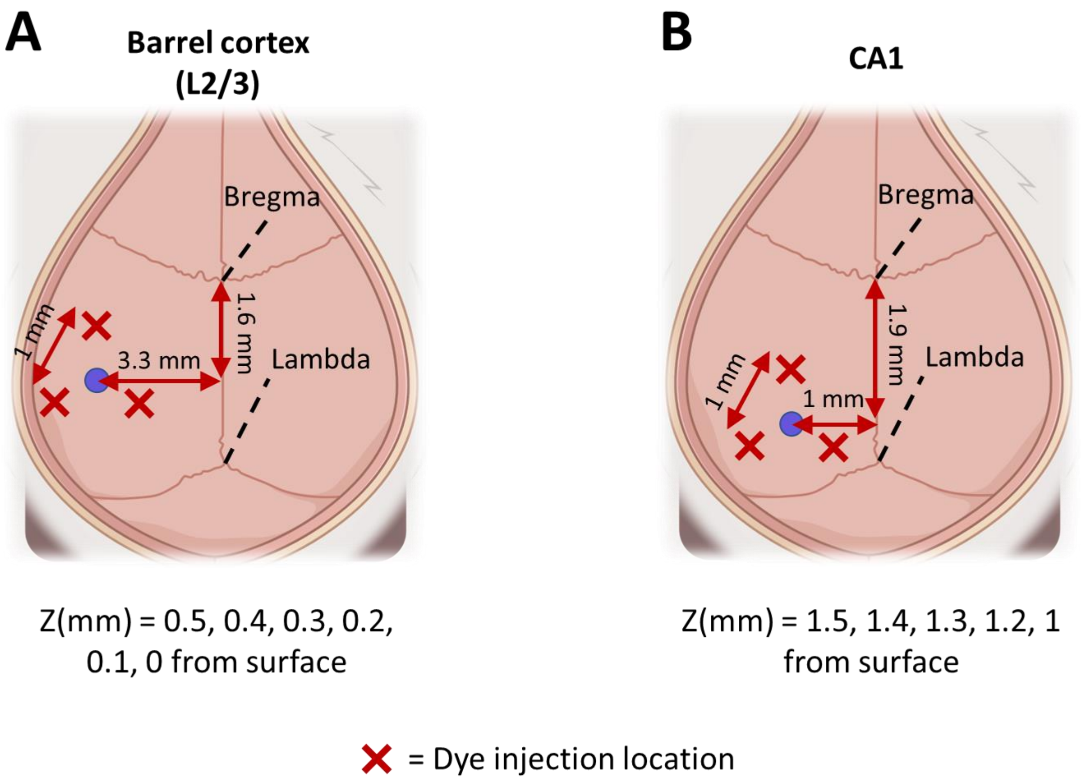


**Fig. S6. Coordinates for *in vivo* dye injections.**

**(A)** Relative coordinates from bregma and lambda for intracortical dye injections to cover layer 2/3 pyramidal neurons in left barrel cortex. **(B)** Relative coordinates from bregma and lambda for intrahippocampal dye injections to cover CA1 pyramidal neurons in left hippocampus.

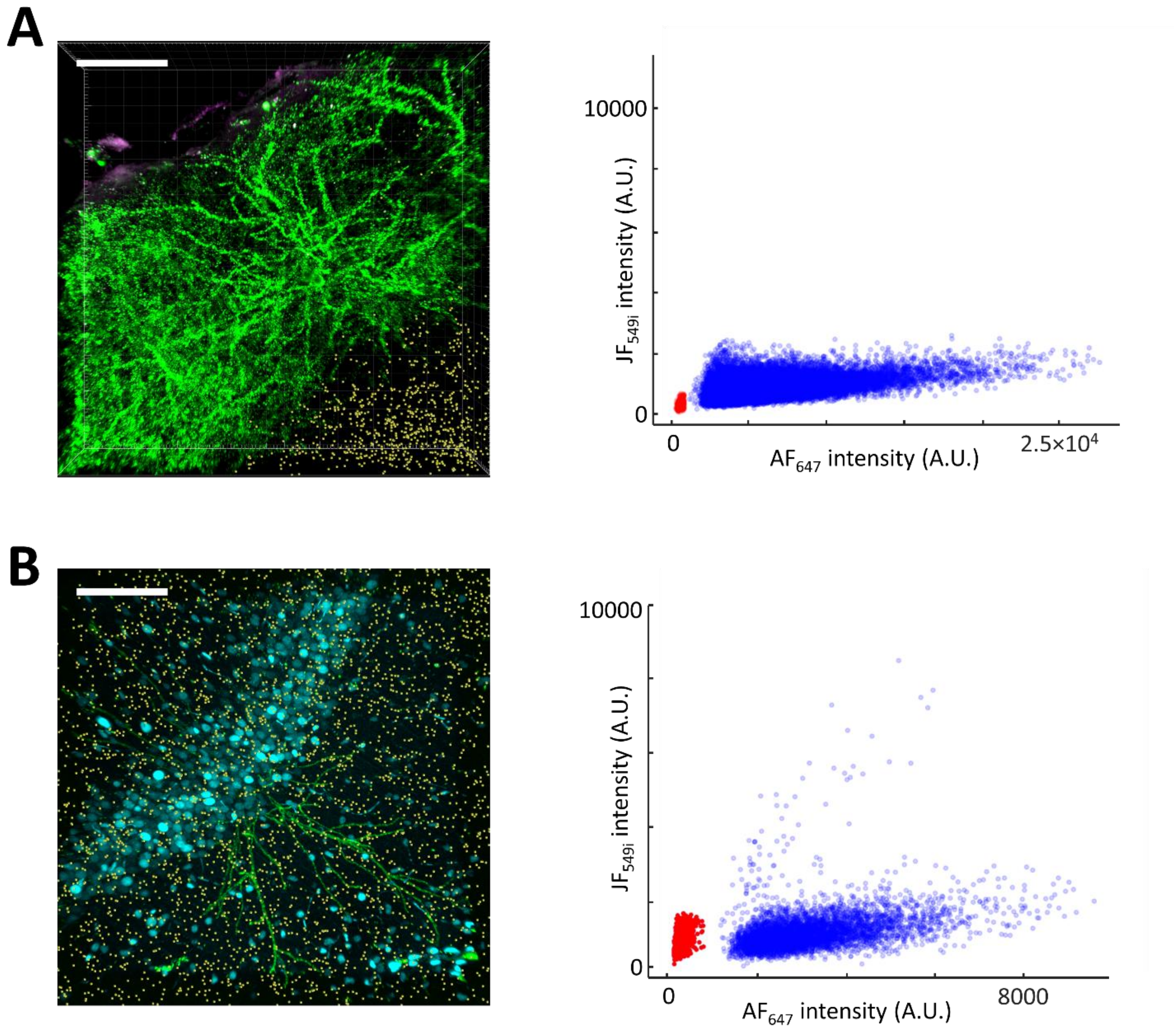


**Fig. S7. Analysis on the background fluorescence.**

**(A)** Background fluorescence analysis in mouse barrel cortex layer 2/3. Left: Spherical regions of interest (yellow) used for background fluorescence measurement on top of Fig. 2G. Right: Scatterplots of spine (blue) and background (red) fluorescence intensities ($n$ = 53,457 spines and 747 background regions). **(B)** Background fluorescence analysis in mouse CA1. Left: Spherical regions of interest (yellow) used for background fluorescence measurement on top of Fig. 3C. Right: Scatterplots of spine (blue) and background (red) fluorescence intensities ($n$ = 5314 spines and 2836 background regions). Scale bars: 200 µm.

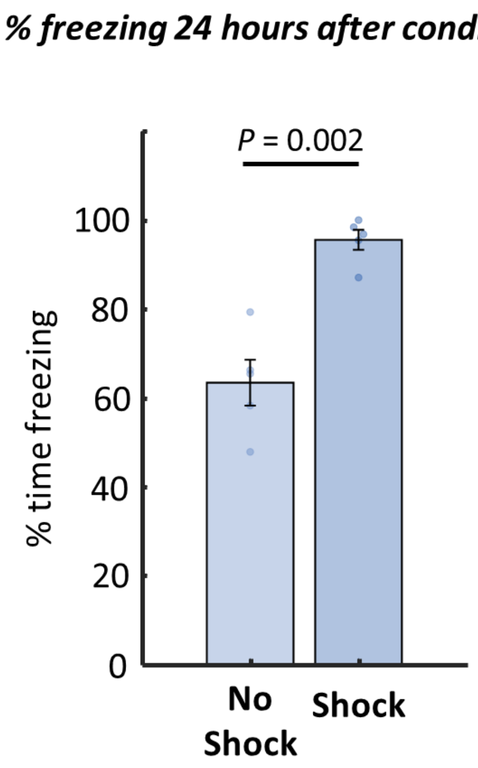


**Fig. S8. Validation of contextual fear conditioning.** The CFC system was validated by measuring the percent of time mouse spent freezing (i.e. immobile) 24 hours after conditioning. The percentage of freezing was measured for the mice that underwent full conditioning (shock) and for the mice exposed to the identical context but that did not receive an aversive stimulus (no shock). Error bars represent mean ± s.e.m. (*n* = 5 mice for each group). Two-sided Wilcoxon rank-sum test.

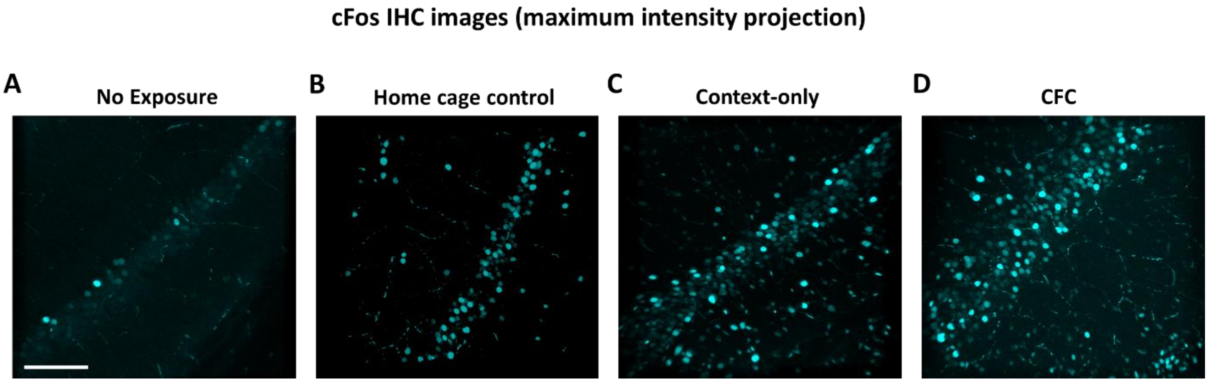


**Fig. S9. cFos IHC images in hippocampus CA1.**

Representative images from CA1 from mice that **(A)** were not exposed to any context (i.e. euthanized after 3 hours of anesthesia), **(B)** were not exposed to the novel context (stayed in their home cages, home cage control), **(C)** were exposed to the novel context but did not receive an aversive foot shock (context-only), and **(D)** underwent CFC. The images are shown in the same contrast scale and are maximum-intensity projections of z-stacks. Scale bars: 200 µm.

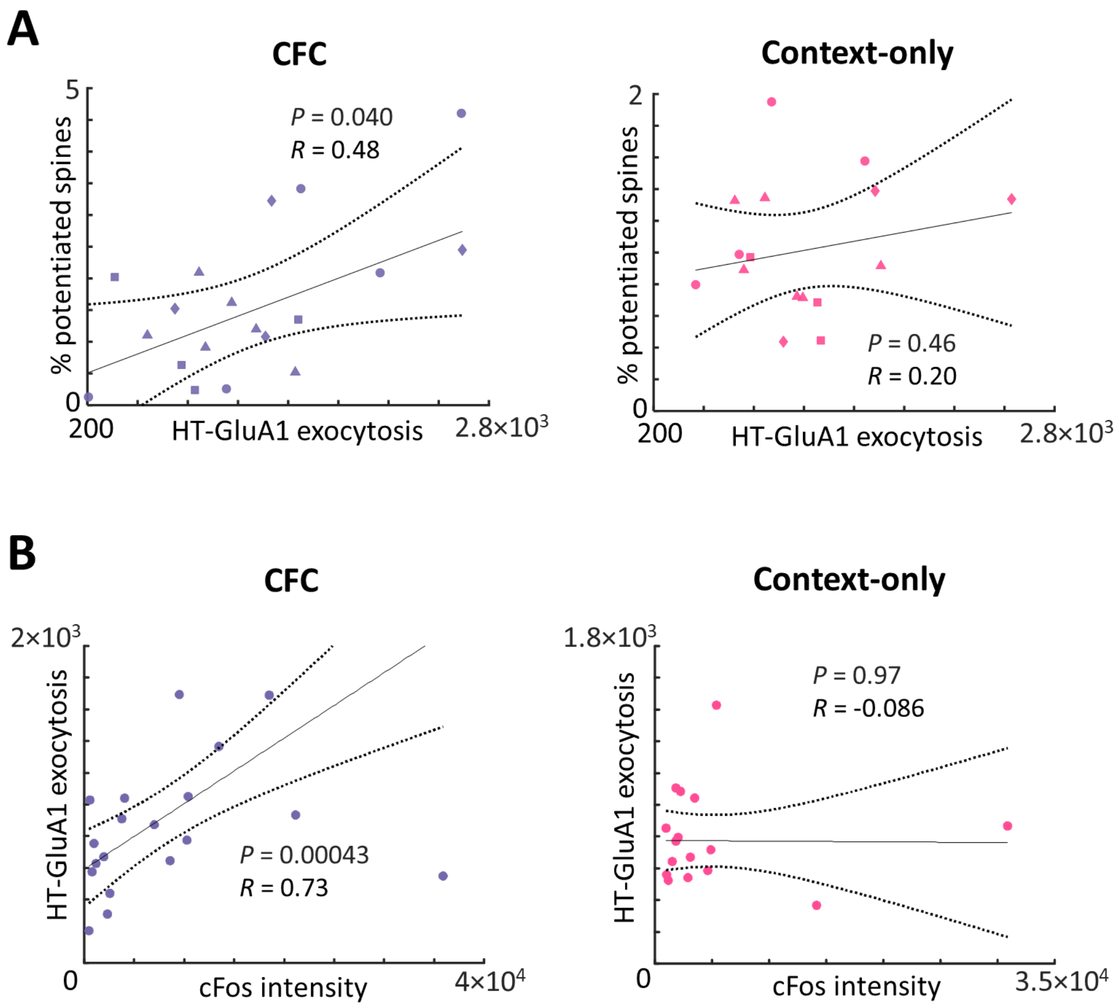


**Fig. S10. Relations among cFos expression, level of AMPAR exocytosis, and percentage of potentiated spines.**

**(A)** Relation of the percentage of potentiated spines to the mean HT-GluA1 exocytosis among potentiated spines (distance of Dye 2 signal above threshold, averaged over above-threshold spines), for (left) CFC and (right) context only control. **(B)** Relation between the mean HT-GluA1 exocytosis among potentiated spines and the corresponding cFos intensity. CFC: $n$ = 19 cells from 4 animals; context-only: $n$ = 16 cells from 4 animals. $R$, Pearson's linear correlation coefficient, $P$ value from two-sided Student's t-test. Distinct animals represented by different shape symbols.

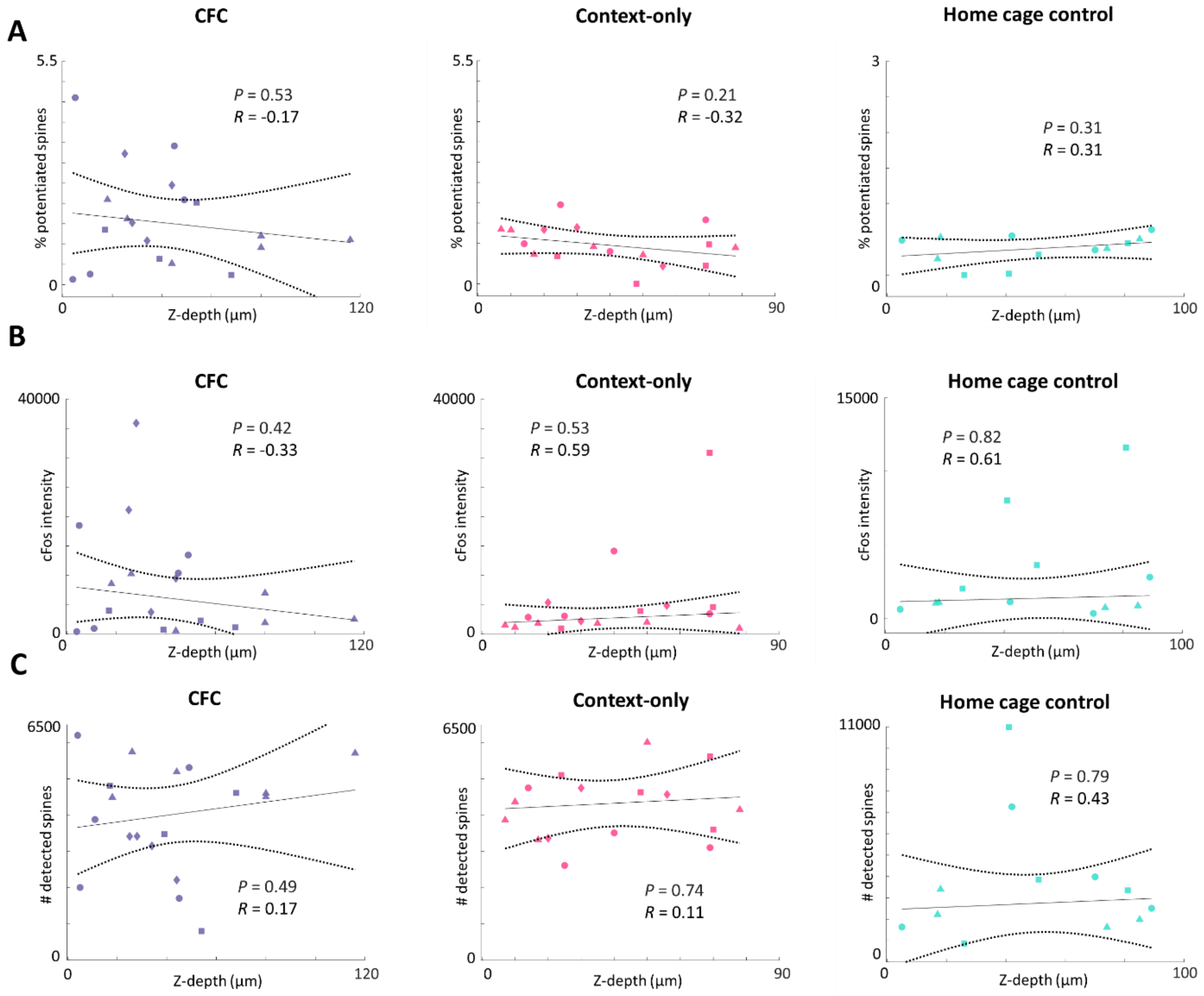


**Fig. S11. Tests for depth-dependent sources of spurious correlation between measures of synaptic plasticity and cFos.**

**(A)** Relation between the percentage of potentiated spine and the z-coordinate of the center of soma of the corresponding neuron for (left) CFC, (middle) context-only control, and (right) home cage control. **(B)** Relation between cFos intensity and the z-coordinate of the center of soma of the corresponding neuron for (left) CFC, (middle) context-only control, and (right) home cage control. **(C)** Relation between the total number of identified spines and the z-coordinate of the center of soma of the corresponding neuron for (left) CFC, (middle) context-only control, and (right) home cage control. CFC: $n$ = 19 cells from 4 animals; context-only: $n$ = 17 cells from 4 animals; home cage control: $n$ = 12 cells from 3 animals. $R$, Pearson's linear correlation coefficient, $P$ value from two-sided Student's t-test. Distinct animals represented by different shape symbols.

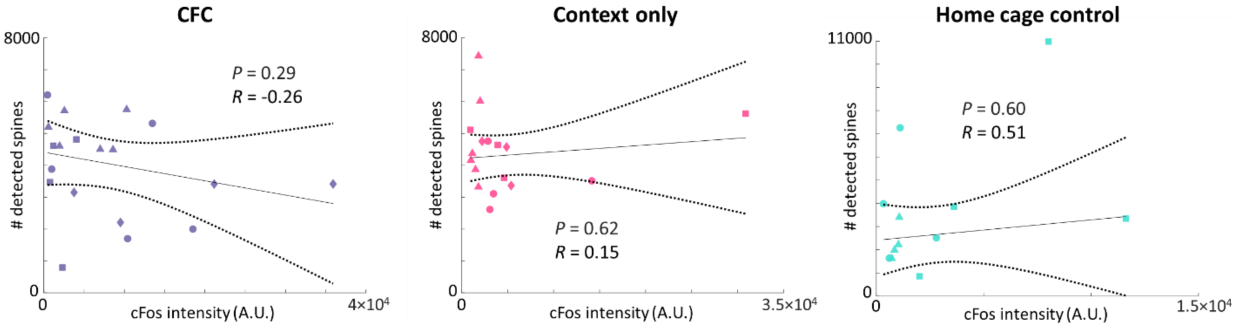


**Fig. S12.** Relation between the total number of identified spines and the cFos intensity of the corresponding neuron for (left) CFC, (middle) context-only control, and (right) home cage control. CFC: $n$ = 19 cells from 4 animals; context-only: $n$ = 17 cells from 4 animals; home cage control: $n$ = 12 cells from 3 animals. $R$, Pearson's linear correlation coefficient, $P$ value from two-sided Student's t-test. Distinct animals represented by different shape symbols.

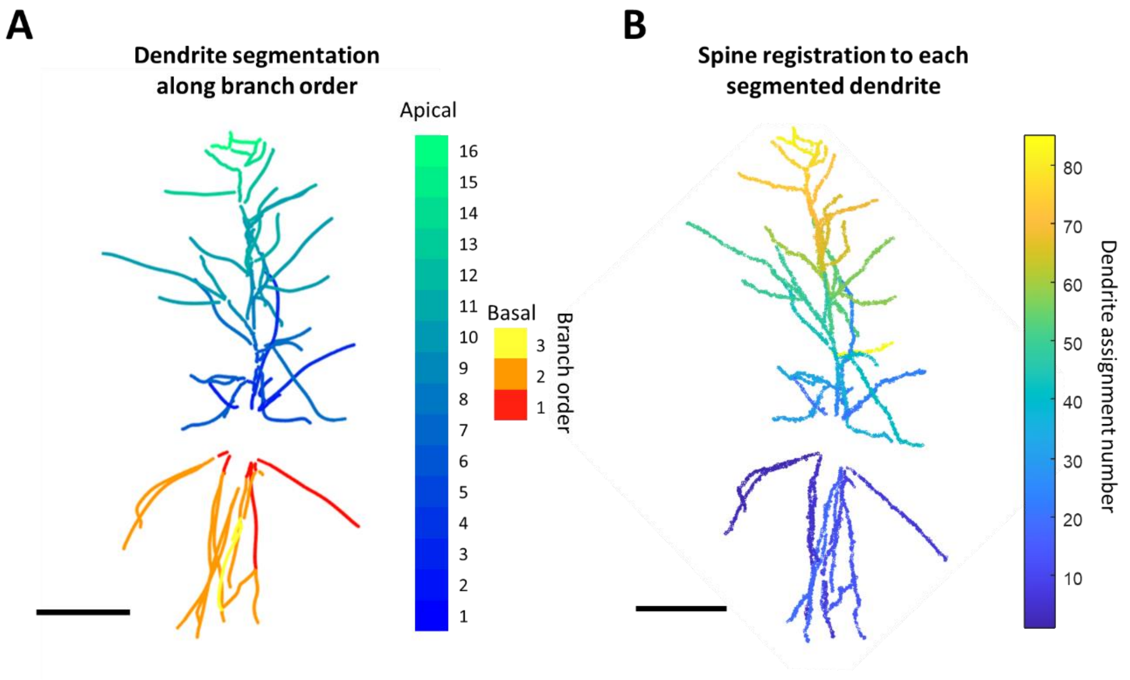


**Fig. S13. Dendrite segmentation and spine registration**

**(A)** Representative neuron with dendrites segmented and colored by their branch order. **(B)** Same neuron with spines registered to the nearest segmented dendrites. Scale bars: 100 μm.

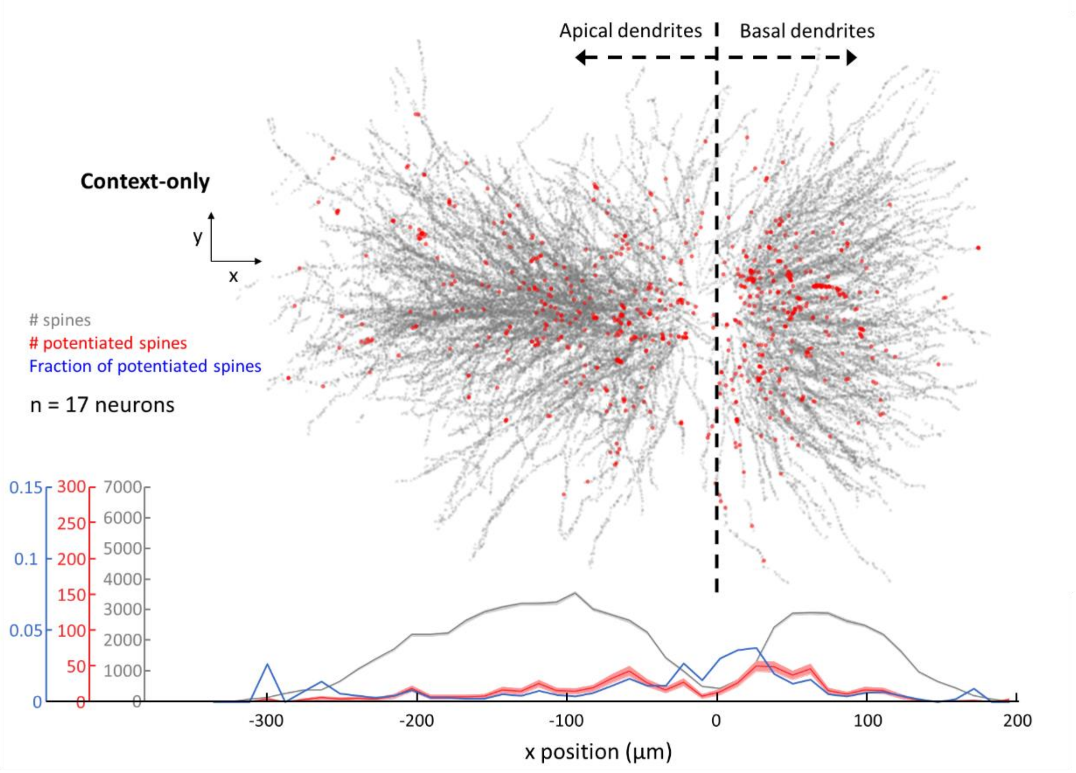


**Fig. S14.** Distribution of potentiated spines as a function of projected distance from the *stratum pyramidale*. (Top) Overlapped images of all identified spines (grey) and spines with high HT-GluA1 exocytosis (red) from the context-only control group. (Bottom) Total number of identified spines (grey), number of spines with high HT-GluA1 exocytosis (red), and fraction of potentiated spines (blue) plotted against the projected distance from the *stratum pyramidale* (x-axis from the top panel). Error bars represent count ± sqrt(count).

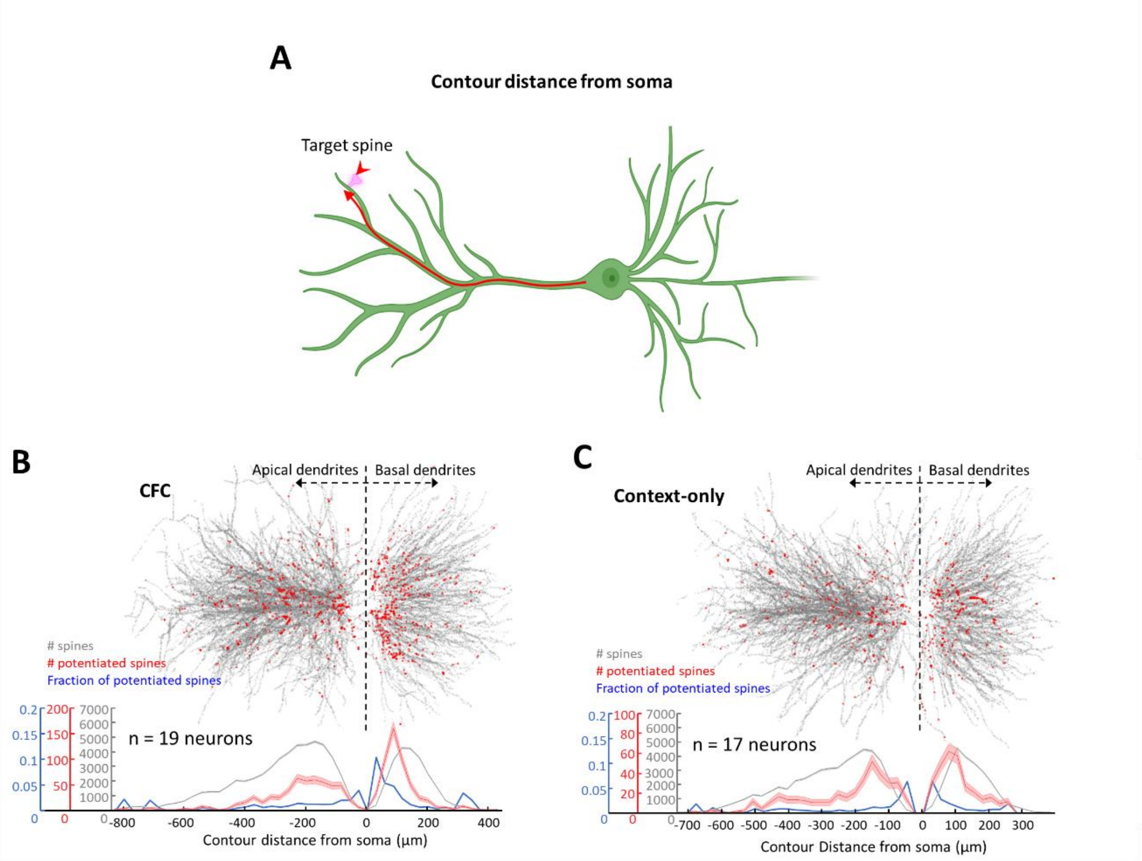


**Fig. S15. Distribution of potentiated spines vs. contour distance from the *stratum pyramidale*.**

**(A)** Schematic diagram showing the contour distance of a neuron's spine from its soma. This is the distance along the dendrite backbone. **(B, C)** Same data as in Fig. 4A and Fig. S14, but plotted vs contour distance instead of *x*.

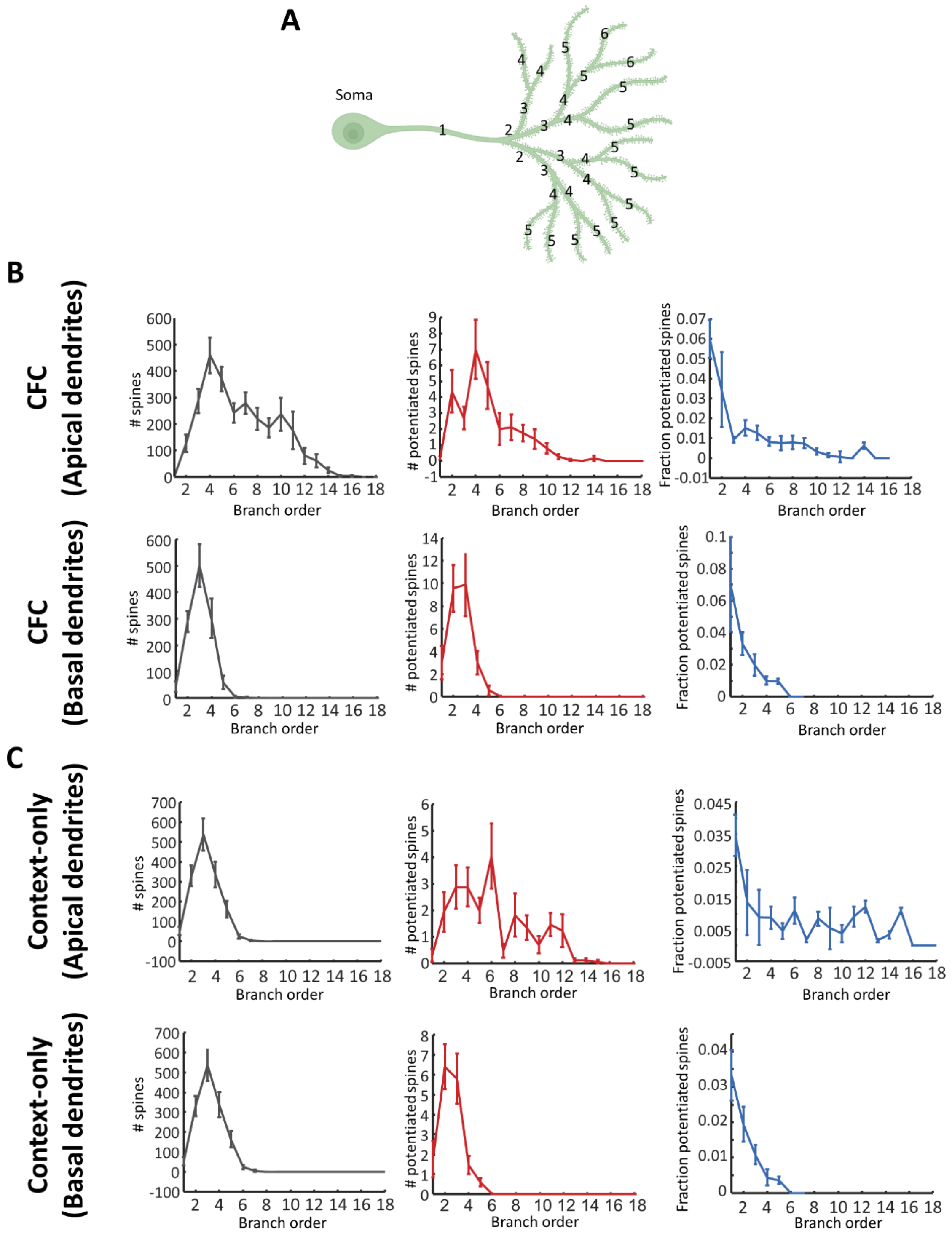


**Fig. S16. Distribution of potentiated spines vs. dendritic branch order.**

**(A)** Schematic drawing of a neuron with dendrites numbered by their branch order. **(B, C)** Total number of identified spines (left), number of potentiated spines (middle), fraction of potentiated spines (right) vs corresponding dendrites' branch order from (top) apical dendrites or (bottom) basal dendrites in **(B)** CFC group and **(C)** context-only controls. CFC: *n* = 19 neurons; context-only: *n* = 17 neurons. Data are represented as mean ± s.e.m.

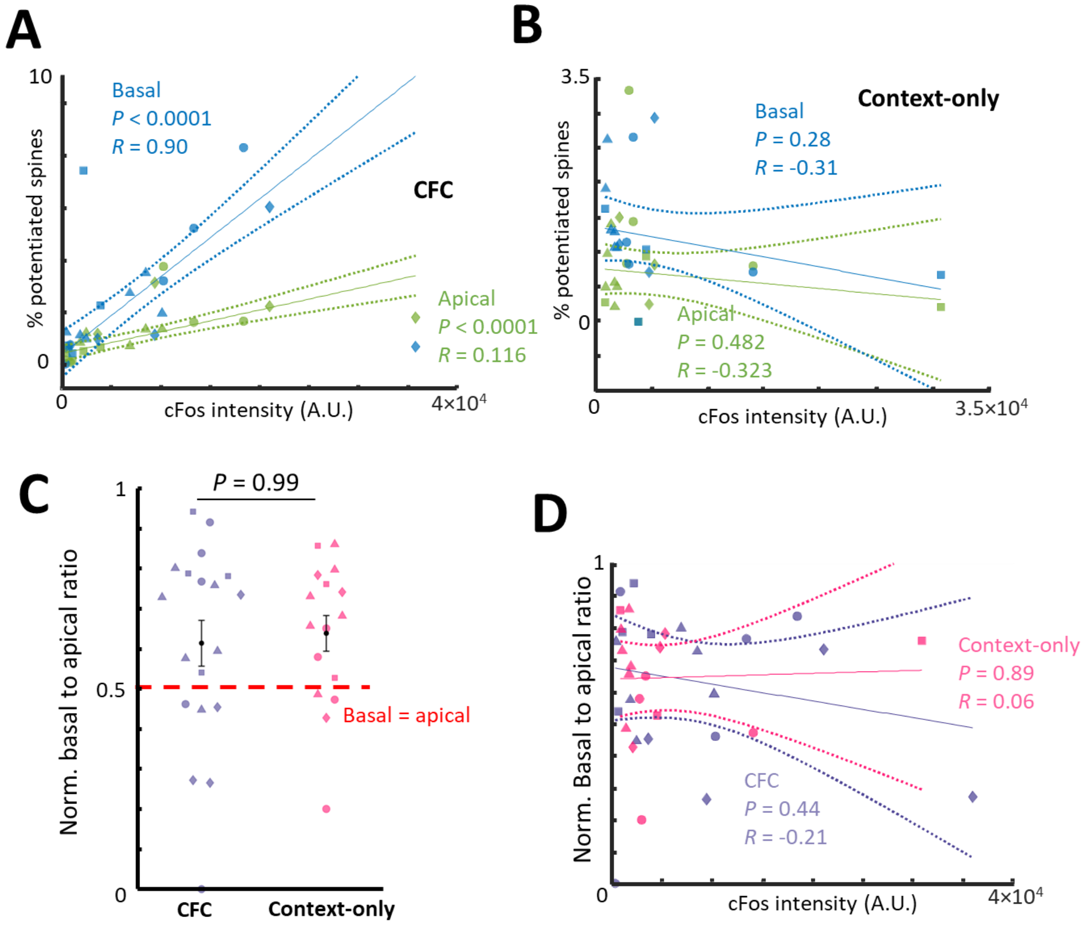


**Fig. S17. Comparison of the fraction of potentiated spines in basal vs. apical dendrites.**

Fraction of potentiated spines for each neuron vs. cFos intensity for **(A)** CFC ($n$ = 19 neurons) and **(B)** context-only control ($n$ = 17 neurons). Spine fraction was separately evaluated for basal and apical dendrites. **(C)** Normalized basal to apical ratio of percent potentiated spines. Normalized ratio: basal percent potentiated spines / (basal percent potentiated spines + apical percent potentiated spines). 13 of 19 neurons in the CFC group and 12 of 16 neurons in the context-only group had higher basal than apical fraction of potentiated spines. There was no significant difference between the CFC and context-only groups in the ratio of basal to apical potentiated spines (CFC: 0.61 ± 0.06, mean ± s.e.m.; context-only: 0.64 ± 0.04, mean ± s.e.m.). Two-sided Wilcoxon rank-sum test. **(D)** Basal to apical ratio of percent potentiated spines vs. cFos intensity. The basal to apical ratio was not correlated with the corresponding cFos levels. $R$, Pearson's linear correlation coefficient, $P$ value from two-sided Student's t-test. Distinct mice represented by different shape symbols.

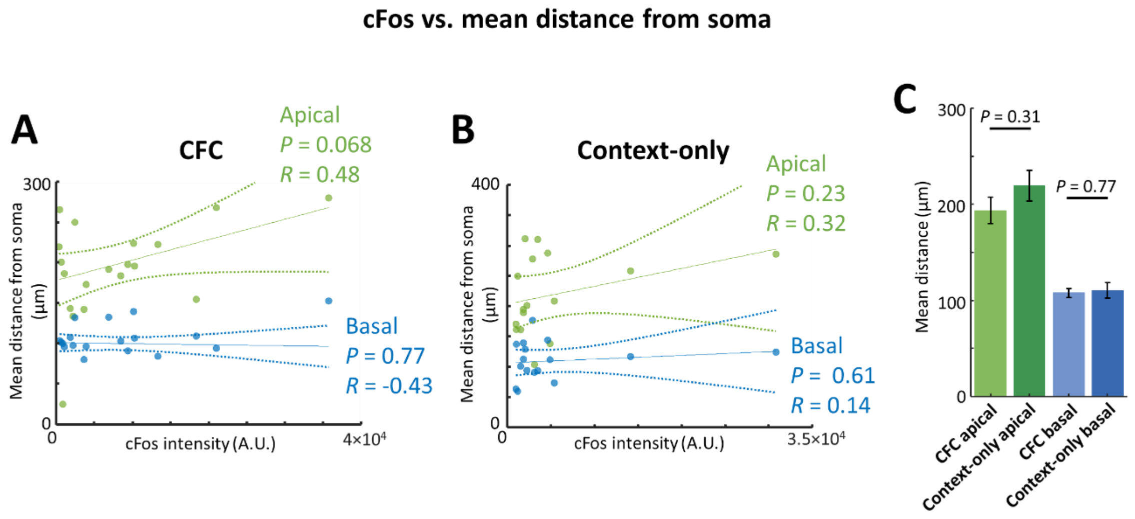


**Fig. S18. Mean contour distance from soma to potentiated spines vs. the corresponding cFos intensity.**

Mean contour distance from soma to potentiated spines vs. corresponding cFos intensity for **(A)** CFC group and **(B)** context-only control. The mean distance was separately evaluated for spines in basal and apical dendrites. *R*, Pearson's linear correlation coefficient, *P* value from two-sided Student's t-test. **(C)** Mean contour distance from soma to potentiated spines in CFC and context-only group. Data are represented as mean ± s.e.m. Two-sided Wilcoxon rank-sum test. CFC: $n$ = 19 neurons; context-only: $n$ = 16 neurons.

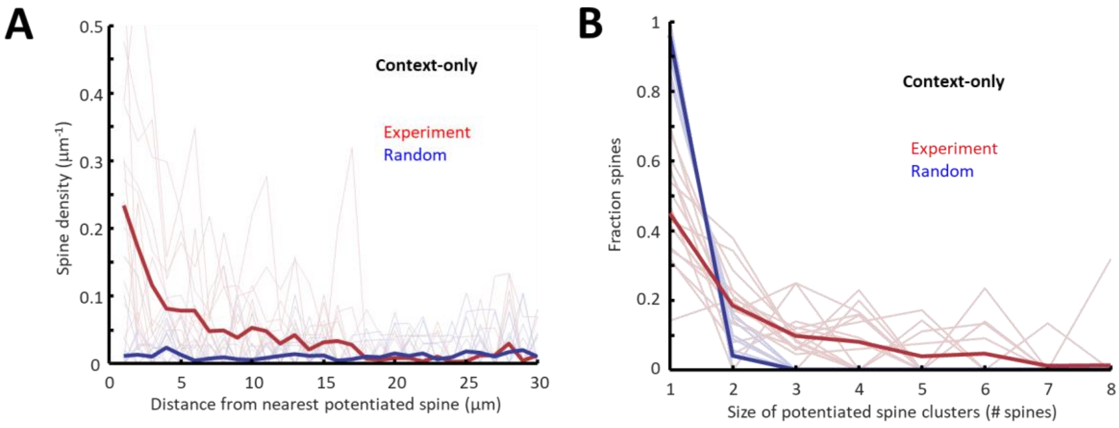


**Fig. S19. Clustering of potentiated spines in context-only control.**

**(A)** Density profile of potentiated spines as a function of distance from the nearest potentiated spine. The single-cell profiles are plotted with light colors. Random: simulation where the same number of potentiated spines are distributed randomly and independently among all detected spines. **(B)** Fraction of potentiated spine clusters of different sizes from the context-only group (n = 16 neurons). The single-cell profiles are plotted with light colors. Random defined as in **(A)**.
